# Protein Kinase Cγ Mutations Drive Spinocerebellar Ataxia Type 14 by Impairing Autoinhibition

**DOI:** 10.1101/2021.06.24.449810

**Authors:** Caila A. Pilo, Timothy R. Baffi, Alexandr P. Kornev, Maya T. Kunkel, Mario Malfavon, Dong-Hui Chen, Liang-Chin Huang, Cheryl Longman, Natarajan Kannan, Wendy Raskind, David J. Gonzalez, Susan S. Taylor, George Gorrie, Alexandra C. Newton

## Abstract

Spinocerebellar ataxia type 14 (SCA14) is a neurodegenerative disease caused by germline variants in the diacylglycerol (DG)/Ca^2+^-regulated protein kinase C gamma (PKCγ), leading to Purkinje cell degeneration and progressive cerebellar dysfunction. The majority of the approximately 50 identified variants cluster to the DG-sensing C1 domains. Here, we use a FRET-based activity reporter to show that ataxia-associated PKCγ mutations enhance basal activity by compromising autoinhibition. Although impaired autoinhibition generally leads to PKC degradation, the C1 domain mutations protect PKCγ from phorbol ester-induced downregulation. Furthermore, it is the degree of disrupted autoinhibition, not changes in the amplitude of agonist-stimulated activity, that correlate with disease severity. This enhanced basal signaling rewires the brain phosphoproteome, as assessed by phosphoproteomic analysis of cerebella from mice expressing a human PKCγ transgene harboring a SCA14 C1 domain mutation, H101Y. Validating that the pathology arises from disrupted autoinhibition, we show that the degree of impaired autoinhibition correlates inversely with age of disease onset in patients: mutations that cause high basal activity are associated with early onset, whereas those that only modestly increase basal activity, including a previously undescribed variant, D115Y, are associated with later onset. Molecular modeling indicates that almost all SCA14 variants that are not in the C1 domains are at interfaces with the C1B domain, and bioinformatics analysis reveals that variants in the C1B domain are under-represented in cancer. Thus, clustering of SCA14 variants to the C1B domain provides a unique mechanism to enhance PKCγ basal activity while protecting the enzyme from downregulation, deregulating the cerebellar phosphoproteome.

**One Sentence Summary:** SCA14 driver mutations in PKCγ impair autoinhibition, with defect correlating inversely with age of disease onset.

## Introduction

Conventional protein kinase C (PKC) isozymes play key roles in normal brain physiology, where they regulate neuronal functions such as synapse morphology, receptor turnover, and cytoskeletal integrity (*1*). These isozymes are transiently and reversibly activated by Ca^2+^ and diacylglycerol (DG), the two second messenger products of receptor-mediated hydrolysis of phosphatidylinositol-4,5-bisphosphate (PIP2) (*2*). Tight control of not only activity, but also steady-state protein levels, is necessary for cellular homeostasis, with deregulation of either resulting in pathophysiology. In general, loss-of-function of conventional PKC isozymes is associated with cancer whereas gain-of-function correlates with neurodegenerative diseases (*3, 4*). Thus, whereas reduced protein levels and activity of conventional PKC isozymes are associated with poorer patient survival in cancers such as colon and pancreatic cancer, enhanced activity of the conventional PKCα is associated with Alzheimer’s disease (*5, 6*).

Spinocerebellar ataxias (SCAs) are a group of over 40 autosomal dominant neurodegenerative diseases characterized by Purkinje cell degeneration and cerebellar dysfunction, resulting in progressive ataxia and loss of motor coordination and control (*7*). Each subtype of SCA is caused by germline variants in distinct genes. A majority of these genes encode proteins that regulate Ca^2+^ homeostasis, including the IP3 receptor, IP3R1 (SCA 15, 16 and 29), ataxins 2 and 3, which regulate IP3R1 function (SCA2 and 3, respectively) (*8, 9*), the cation channel TRPC3 (SCA41) (*10*), and mGluR1 which couples to phospholipase C (SCA44) (*11*). Spinocerebellar ataxia type 14 (SCA14) is caused by missense variants in PKCγ (*12*), a conventional PKC isozyme whose expression is restricted to neurons, particularly Purkinje cells (*13, 14*). Given that Ca^2+^ is an important activator of PKC, one intriguing theory is that enhanced PKCγ activity is not only central to SCA14 pathology, but is also at the epicenter of many other types of SCA. Thus, understanding how SCA14-associated variants deregulate the function of PKCγ has strong potential clinical relevance.

Exquisite regulation of the spatiotemporal dynamics of conventional PKC signaling ensures that these enzymes are only activated for a specific time, at defined locations, and in response to appropriate stimuli. In the absence of specific stimuli, these enzymes are maintained in an autoinhibited conformation by an N-terminal regulatory moiety that constrains the catalytic activity of the C-terminal kinase domain (*15*). Specifically, an autoinhibitory pseudosubstrate segment occupies the substrate-binding cavity to maintain the enzyme in an inactive conformation. Additionally, multiple interactions of the kinase domain with modules in the regulatory moiety secure the pseudosubstrate in place to prevent aberrant basal signaling. These modules are the DG-sensing C1A and C1B domains and Ca^2+^-sensing C2 domain, which pack against the kinase domain to maintain it in an autoinhibited conformation until the relevant second messengers are generated (*16*). Release of the pseudosubstrate occurs upon generation of the appropriate second messengers. Specifically, following phospholipase C-catalyzed hydrolysis of PIP2, Ca^2+^ binds to the C2 domain causing it to translocate to the plasma membrane where it is anchored by interaction of a basic surface with PIP2 (*17*). At the membrane, the C1B domain engages its membrane-embedded allosteric activator, DG, resulting in release of the pseudosubstrate from the active site, allowing PKC to phosphorylate its substrates (*18*). This process is readily reversible upon decay of the second messengers, and thus normal PKC activity is transient. Before PKC can adopt an autoinhibited but signaling-competent conformation, newly synthesized enzyme must be processed by a series of ordered phosphorylations in the kinase domain. In particular, phosphorylation at a residue termed the hydrophobic motif is required for PKC to adopt the autoinhibited conformation (*19*). Aberrant PKC that is not properly autoinhibited is dephosphorylated by the phosphatase PHLPP, ubiquitinated, and degraded by a proteasomal pathway (*19*). This quality control mechanism ensures that only properly autoinhibited PKC accumulates in the cell. For example, cancer-associated variants that prevent autoinhibition of PKC are paradoxically loss-of-function because the mutant protein is degraded by this quality control pathway (*19*). Thus, autoinhibited PKC is stable and prolonged activation renders PKC sensitive to dephosphorylation and degradation. In this regard, phorbol esters which bind PKC with high affinity and are not readily metabolized cause the acute activation but long-term downregulation of PKC.

Since the original discovery of germline variants in PKCγ by Raskind and colleagues that defined SCA14 (*12, 20, 21*), approximately 50 variants across all domains of PKCγ have now been identified in SCA14 (*22–25*). Mouse model studies by Kapfhammer and colleagues have established that a single SCA14-associated point mutation in PKCγ is sufficient to drive pathophysiology characteristic of the human disease, including Purkinje cell degeneration and motor deficits (*26, 27*). Cellular studies by several groups have addressed the mechanism by which cerebellar degeneration in SCA14 may precipitate. Schrenk *et al.* have shown that stimulation of PKC in mouse cerebellar slices by treatment with phorbol esters leads to a decrease in Purkinje cell dendrites, whereas inhibition of PKC leads to hyper-arborization, suggesting a causative role for enhanced PKC activity in Purkinje cell degeneration (*28*). Others have observed that PKC inhibition prevents Purkinje cell death (*29*). Verbeek and colleagues have also identified a role for altered PKCγ activity in SCA14, showing in some cases that SCA14-associated PKCγ mutations lead to unmasking of the C1 domains to enhance ‘openness’ and thus membrane accessibility of PKCγ, but concluded that these mutants have lower kinase activity (*30, 31*). A sizable body of work has also focused on the role of PKCγ aggregation in SCA14. Notably, Saito and colleagues have shown that in both overexpression and *in vitro* systems, wild-type and mutant PKCγ form amyloid-like fibrils and aggregates that lead to cell death, which can be decreased by pharmacological induction of heat shock proteins (*32, 33*). Other studies have also demonstrated the presence of such aggregates in iPSCs from SCA14 patients or primary culture mouse Purkinje cells (*23, 34*). However, the precise biochemical mechanisms in which SCA14 mutations alter PKCγ function to ultimately drive neurodegeneration in SCA14 is still unknown.

Here, we used our genetically-encoded biosensor for PKC activity, coupled with biochemical, molecular modeling, and bioinformatics approaches, to address the mechanism by which SCA14 mutations affect PKCγ function. Our studies reveal that SCA14-associated mutations in every segment or domain of PKCγ (pseudosubstrate, C1A, C1B, C2, kinase) produce the same defect: impaired autoinhibition leading to increased basal activity. Furthermore, we show that SCA14-associated PKCγ mutations in the C1A and C1B domains, mutational hotspots for the disease, render PKCγ insensitive to phorbol ester-mediated downregulation, an effect also observed by deletion of either domain. Specifically, mutation in (or deletion of) the C1A domain prevented dephosphorylation, the first step in downregulation, and mutations in (or deletion of) the C1B domain permitted dephosphorylation but prevented the next step, protein degradation. Thus, C1A and C1B domain mutations provide unique mechanisms to deregulate PKC without subjecting it to degradation. Focusing on one mutation in the C1A domain, ΔF48, we show that deletion of this single residue (or the entire C1A domain) not only reduces autoinhibition resulting in high basal activity but also uncouples communication between the pseudosubstrate and the kinase domain to trap this PKC in an unresponsive but slightly ‘open’ state. Structural analyses reveal that most SCA14 mutations are either in the C1 domains or at common interfaces with the C1 domains. Furthermore, bioinformatics analyses reveal that mutations in the C1 domains are relatively under-represented in cancer, a disease where conventional PKC function is generally lost. This is consistent with our findings that mutations in these domains will enhance, not suppress, PKC activity. Validating altered signaling in a physiological context, phosphoproteomic analysis of cerebella from mice expressing a human bacterial artificial chromosome (BAC) WT or H101Y PKCγ transgene reveals significant alteration in the phosphorylation of components related to cytoskeletal organization and neuronal development. Lastly, compilation of the age of SCA14 onset for C1 domain mutants revealed that the magnitude of the biochemical defect (reduced autoinhibition) inversely correlated with age of SCA14 onset. Taken together, our results reveal that sustained ‘leaky’ activity of PKCγ, by mechanisms that protect it from degradation, alters the cerebellar phosphoproteome to drive SCA14 pathology.

## Results

### PKCγ pathogenic variants cause spinocerebellar ataxia type 14

SCA14 is caused by germline variants in PKCγ, of which over 50 unique variants have been identified (Fig. 1A) (*22–24*). Although these variants occur in every domain of the kinase, the majority cluster to the C1 domains, particularly the C1B domain. This small globular DG-binding domain coordinates two Zn^2+^ ions via invariant histidine and cysteine residues (Fig. 1A, residues of motif in red). Mutation of any of the Zn^2+^-coordinating residues abolishes or severely impairs phorbol ester binding (*35*). The SCA14 C1B variants occur with the highest frequency at residues within the zinc finger motif, suggesting that these mutants may affect ligand binding to C1B, and thus, proper regulation of kinase activity. Here, we also report on a previously undescribed variant, D115Y, identified by whole-genome sequencing of a patient who was diagnosed with ataxia. This patient’s mother came from a large family with 6 out of 12 siblings diagnosed with ataxia, consistent with the autosomal dominant nature of the disease. Magnetic resonance imaging (MRI) on the patient harboring the novel D115Y variant revealed significant cerebellar degeneration when compared with a healthy, age-matched individual (Fig. 1B). Of the subset of this patient’s family who underwent whole-genome sequencing (indicated by black outline), three individuals diagnosed with ataxia harbored the D115Y variant (red), while the one healthy individual sequenced did not harbor this variant (blue) (Fig. 1C), indicating segregation of the variant with the disease.

**Figure 1.**
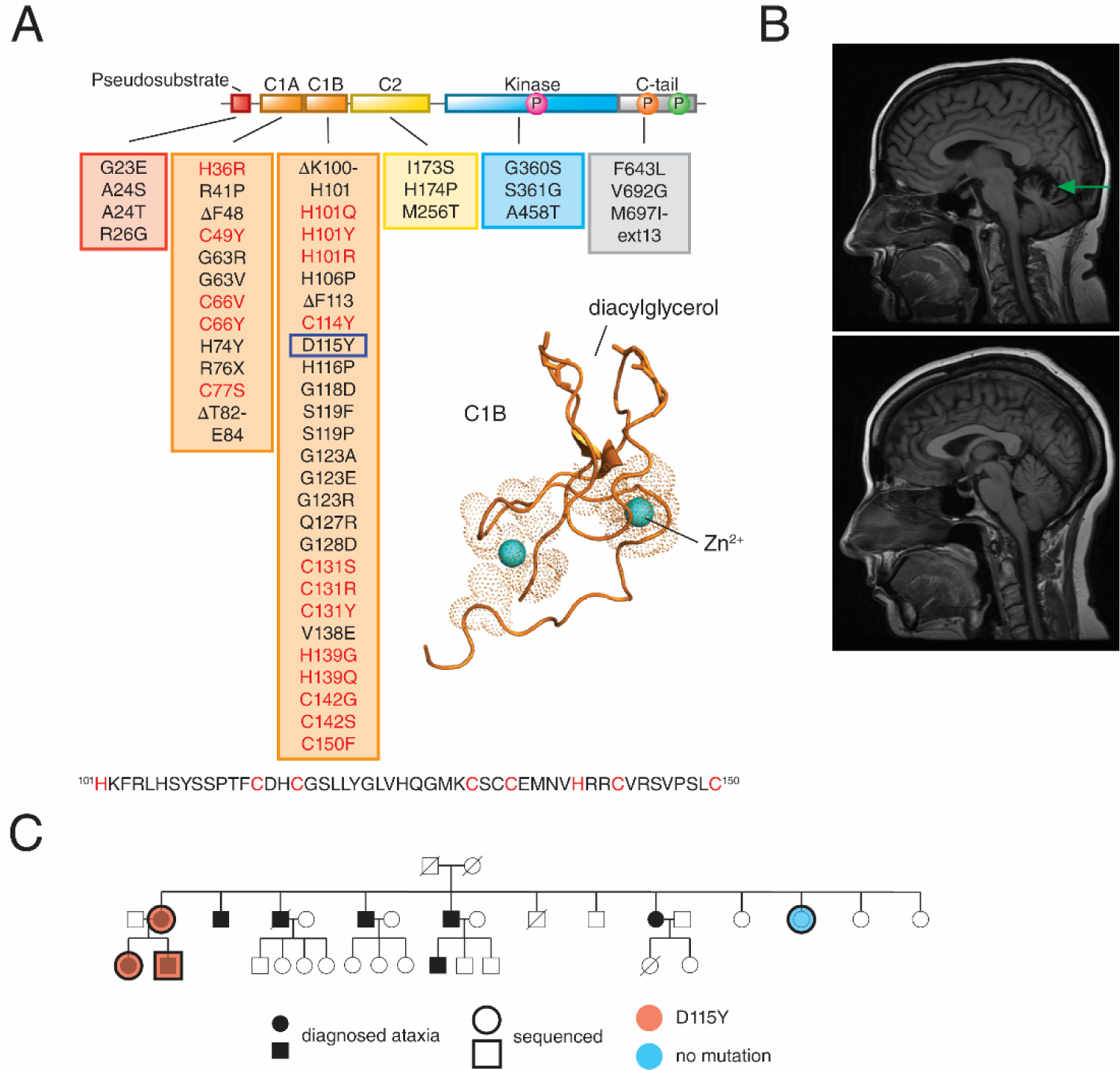
PKCγ in Spinocerebellar Ataxia Type 14. (A) Primary structure of PKCγ with all known SCA14 variants indicated in boxes beneath each domain (*22–25*). Newly identified patient variant (D115Y) indicated with blue box. Previously published crystal structure (*59*) of PKCβII C1B domain shown with Zn^2+^ (cyan spheres) and diacylglycerol binding sites labeled (PDB: 3PFQ). Conserved His and Cys residues of Zn^2+^ finger motif are shown in red in PKCγ primary sequence. (B) MRI of patient with D115Y variant (top) compared to age-matched healthy control (bottom); green arrow indicates cerebellar atrophy. (C) Pedigree of family with PKCγ D115Y variant; black shape-fill indicates family members diagnosed with ataxia, overlaid shapes indicate family members that have been sequenced, blue-filled shape indicates family member with no variant, red-filled shapes indicate family members with D115Y variant.

### SCA14-associated PKCγ mutants display decreased autoinhibition

To assess how SCA14 mutations affect PKCγ function, we first addressed their effect on the basal and agonist-evoked activity of PKCγ in cells using the genetically-encoded FRET-based biosensor, C Kinase Activity Reporter 2 (CKAR2) (*36*). Mutations in each domain were selected for analysis, including the new D115Y mutation in the C1B domain. Additionally, constructs lacking the pseudosubstrate segment (ΔPS) or regulatory domain (ΔC1A, ΔC1B, or ΔC2) were analyzed. COS7 cells co-expressing mCherry-tagged PKCγ constructs and the reporter were sequentially treated with 1] uridine-5′-triphosphate (UTP), which activates purinergic receptors to elevate diacylglycerol (DG) and Ca^2+^, to transiently activate PKC, 2] phorbol 12,13-dibutyrate (PDBu) to maximally activate PKC, and 3] the phosphatase inhibitor Calyculin A to assess maximal phosphorylation of the reporter; traces were normalized to this endpoint. UTP stimulation of cells caused a transient activation of endogenous (grey) and overexpressed wild-type (WT) PKCγ (orange) that was reversed as the enzyme regained the autoinhibited conformation following second messenger decay, as previously reported (*37*) (Fig. 2A). Phorbol ester treatment resulted in nearly maximal phosphorylation of the reporter in cells overexpressing WT PKCγ; endogenous PKC required phosphatase suppression with Calyculin A to observe maximal reporter phosphorylation (Fig. 2A). These kinetics are characteristic of properly autoinhibited PKC (*19*). In contrast, the two SCA14 pseudosubstrate mutants (A24T and R26G) had high basal activity resulting in only modest additional activation by UTP and phorbol esters, approaching the level of deregulated autoinhibition observed upon deletion of the entire pseudosubstrate segment (ΔPS) (Fig. 2A). The C1A SCA14 mutation ΔF48, in which a single residue is deleted (no frameshift) also had high basal activity, but was relatively unresponsive to stimulation with UTP or PDBu (Fig. 2B). This signature of high basal activity and lack of response to agonists was also observed upon deletion of the entire C1A domain (ΔC1A). Mutations in the C1B domain, including the new D115Y, all caused an increase in basal activity but, in contrast to the C1A mutations, did not uncouple responsiveness to UTP and PDBu, similar to the effect observed with C1B domain deletion (ΔC1B) (Fig. 2C). Mutations in the C2 domain, as well as deletion of the entire C2 domain, resulted in slightly enhanced basal activity but reduced response to agonist (Fig. 2D). Lastly, mutations in the kinase domain (S361G) and C-tail (F643L) resulted in both an increase in basal activity and an increase in agonist-evoked activity compared with WT PKCγ. Note that experiments using the previously characterized CKAR1 (*38*), under similar experimental conditions, produced the same qualitative results as CKAR2, although CKAR2 displayed a larger dynamic range (Fig. S1). Every mutant tested exhibited higher basal activity compared to WT, but with varying degrees of deregulation, as revealed by quantitation of the initial FRET ratio of each trace, normalized to that of WT enzyme (Fig. 2F). Thus, SCA14 mutations in every domain of PKCγ consistently display impaired autoinhibition.

**Figure 2.**
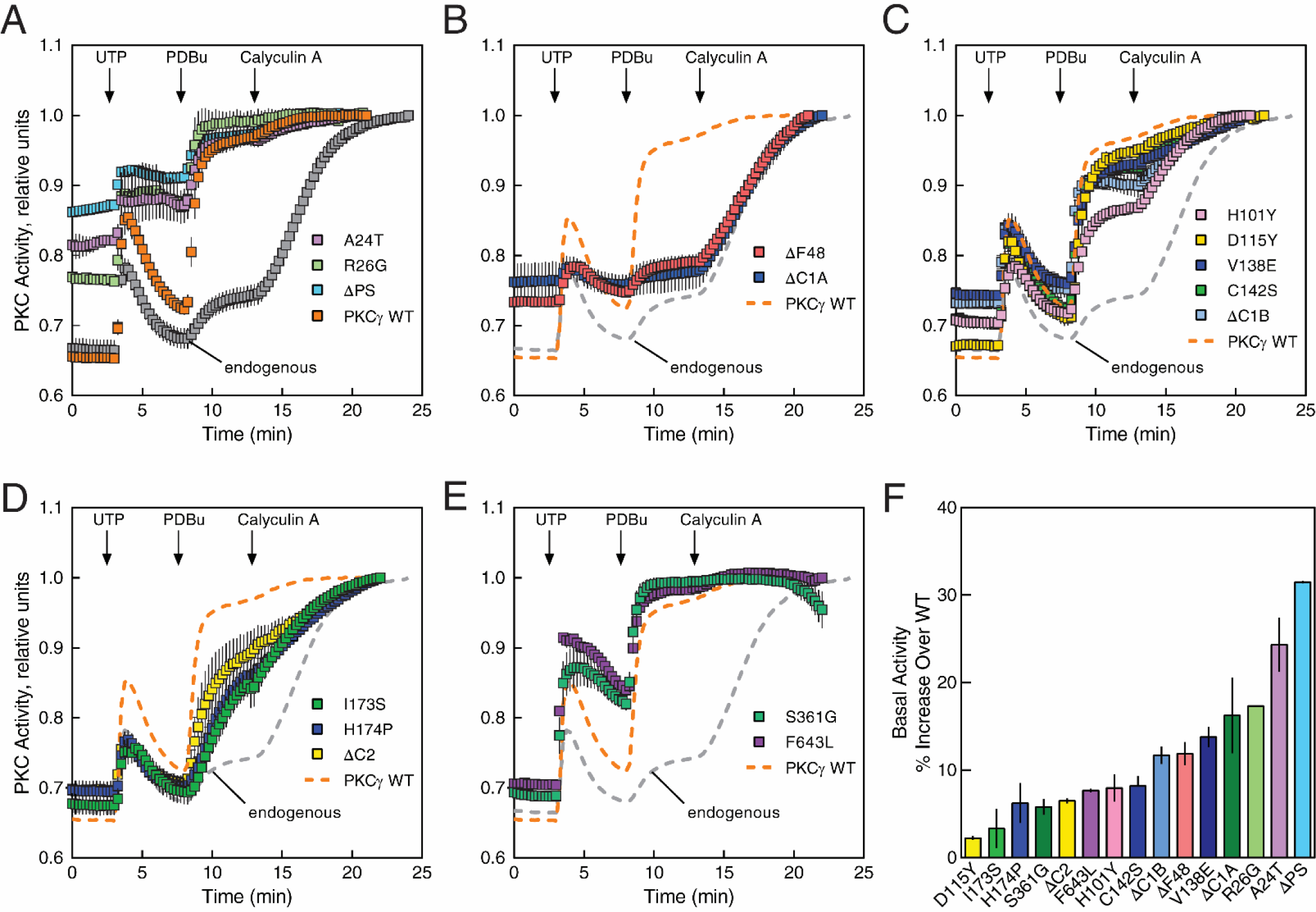
SCA14 mutants exhibit impaired autoinhibition compared to WT PKCγ. (A) COS7 cells were transfected with CKAR2 alone (endogenous) or co-transfected with CKAR2 and mCherry-tagged WT PKCγ (orange), PKCγ lacking a pseudosubstrate (ΔPS; cyan), or the indicated pseudosubstrate SCA14 mutants. PKC activity was monitored by measuring FRET/CFP ratio changes after sequential addition of 100 µM UTP, 200 nM PDBu, and 50 nM Calyculin A at the indicated times. Data were normalized to the endpoint (1.0) and represent mean ± S.E.M. from at least two independent experiments (n ≥ 17 cells per condition). PKCγ WT and endogenous data are reproduced in (**B**)-(**E**) for comparison (dashed lines). (B) As in (**A**), with mCherry-tagged ΔC1A, or ΔF48. Data represent mean ± S.E.M. from at least two independent experiments (n ≥ 16 cells per condition). PKCγ WT and endogenous data are reproduced in for comparison (dashed lines). (C) As in (**A**), with mCherry-tagged ΔC1B, or C1B SCA14 mutants. Data represent mean ± S.E.M. from at least two independent experiments (n ≥ 19 cells per condition). PKCγ WT and endogenous data are reproduced in for comparison (dashed lines). (D) As in (**A**), with mCherry-tagged ΔC2, or C2 SCA14 mutants. Data represent mean ± S.E.M. from at least two independent experiments (n ≥ 11 cells per condition). PKCγ WT and endogenous data are reproduced in for comparison (dashed lines). (E) As in (**A**), with mCherry-tagged SCA14 kinase domain and C-tail mutants. Data represent mean ± S.E.M. from at least two independent experiments (n ≥ 33 cells per condition). PKCγ WT and endogenous data are reproduced in for comparison (dashed lines). (F) Quantification of percent increase in basal activity in (**A**) - (**E**) over WT PKCγ.

PKC with impaired autoinhibition is in a more ‘open’ conformation with its membrane-targeting modules unmasked, resulting in enhanced membrane affinity and faster kinetics of agonist-dependent membrane translocation (*18*). To further characterize how SCA14-associated mutations in the C1 domains affect the ‘openness’ of PKCγ, we examined the translocation of the SCA14 mutants D115Y and ΔF48 compared to WT using a FRET-based translocation assay. Plasma membrane-targeted CFP and YFP-tagged WT, D115Y, ΔC1B, ΔF48, or ΔC1A PKCγ were co-expressed in COS7 cells and the increase in FRET following stimulation of cells with PDBu, a measure of membrane association, was determined (Fig. 3). In response to PDBu, the D115Y mutant associated much more robustly with plasma membrane compared to WT, consistent with unmasking of membrane-targeting modules (Fig. 3A). Furthermore, deletion of the C1B domain (ΔC1B) prevented translocation above WT levels, suggesting that the C1B domain is the predominant binder of plasma membrane-embedded PDBu. On the other hand, deletion of the C1A domain (ΔC1A) enhanced plasma membrane binding, suggesting that the loss of the C1A unmasked the C1B domain to facilitate PDBu binding. ΔF48 translocated with comparable kinetics and magnitude as WT, which could be accounted for by proper masking of its C1B domain (i.e. with normal accessibility to ligand) (Fig. 3B). To further assess enhanced membrane association of the D115Y mutant, mCherry-tagged PKCγ WT and YFP-tagged PKCγ D115Y were co-expressed in COS7 cells, and phorbol ester-stimulated translocation was monitored within the same cells (Fig. 3C). Both WT and D115Y displayed diffuse localization in the cytosol before PDBu treatment. Whereas there was little detectable difference in translocation of the WT PKCγ 4 min following addition of PDBu, D115Y PKCγ displayed enhanced plasma membrane association, which was sustained at 16 minutes post-PDBu addition. Thus, these results are consistent with the D115Y being in a more basally ‘open’ conformation resulting in enhanced association with plasma membrane following phorbol ester treatment.

**Figure 3.**
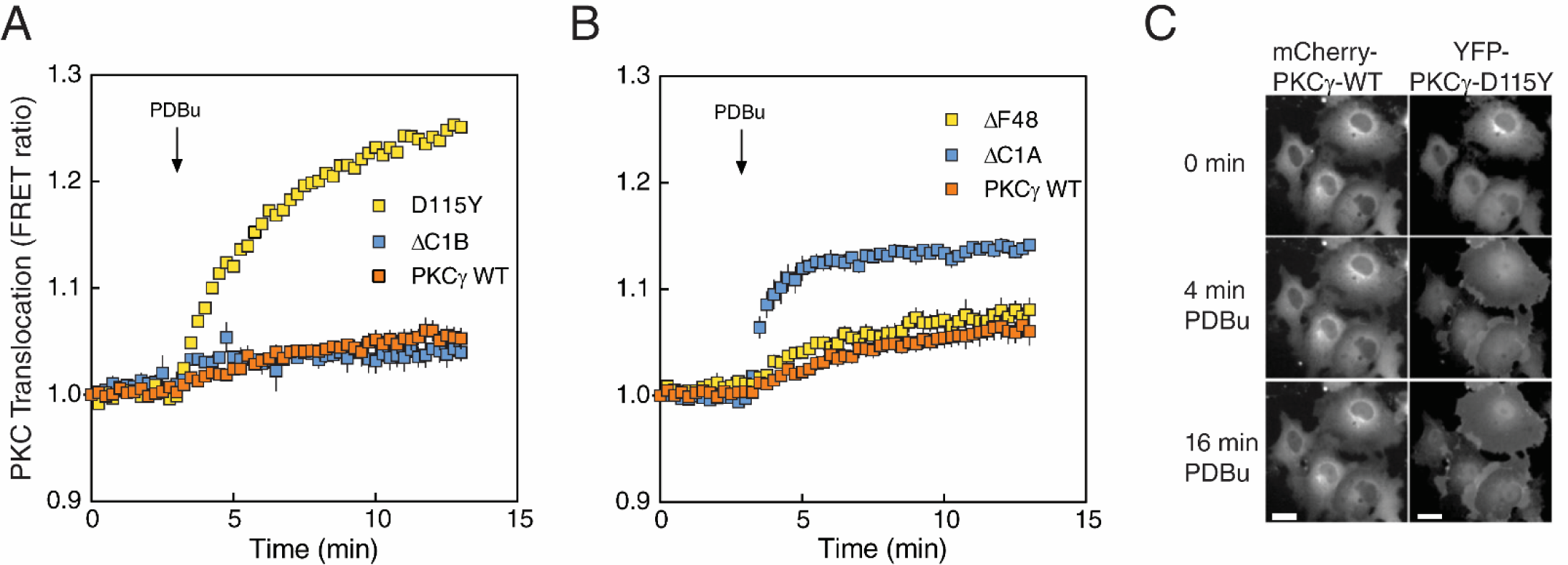
SCA14 mutations affect translocation of PKCγ. (A) COS7 cells were co-transfected with MyrPalm-CFP and YFP-tagged WT PKCγ (orange), PKCγ D115Y (yellow), or PKCγ ΔC1B (blue). Rate of translocation to plasma membrane was monitored by measuring FRET/CFP ratio changes after addition of 200 nM PDBu. Data were normalized to the starting point (1.0) and are representative of two independent experiments (n ≥ 22 cells per condition). (B) As in (**A**), with YFP-tagged ΔF48 or ΔC1A. Data represent mean ± S.E.M. from at least three independent experiments (n ≥ 23 cells per condition). (C) COS7 cells were co-transfected with mCherry-tagged WT PKCγ and YFP-tagged PKCγ D115Y. Localization of mCherry-PKCγ (WT) (left) and YFP-PKCγ-D115Y (right) (same cells) under basal conditions and after addition of 200 nM PDBu was observed by fluorescence microscopy. Images are representative of three independent experiments. Scale bar = 20μm.

### SCA14 mutants evade phorbol ester-mediated degradation, yet display higher turnover

Because reduced autoinhibition of PKC renders the constitutive phosphorylation sites within the kinase domain and C-tail highly phosphatase labile, we examined the phosphorylation state of the basally active SCA14 mutants. Phosphorylation of HA-tagged PKCγ WT, the indicated SCA14 mutants, ΔC1A, or ΔC1B overexpressed in COS7 cells was assessed by monitoring the phosphorylation-induced mobility shift that accompanies phosphorylation of the two C-terminal sites (*39*) or using phospho-specific antibodies to the activation loop (pThr^514^), the turn motif (pThr^655^), and the hydrophobic motif (pThr^674^) by Western blot (Fig. 4A). WT PKCγ migrated predominantly as a slower mobility species (phosphorylated); this slower mobility species was detected with each of the phospho-specific antibodies. In contrast, the ΔC1B migrated as a single species and was not phosphorylated at any of the processing sites (note that for the activation loop (pThr^514^) blot, the band present represents endogenous PKC). Each SCA14 mutant had reduced phosphorylation compared to WT as assessed by the ratio of upper (phosphorylated) to lower (unphosphorylated) bands, with D115Y having the smallest defect and the ΔF48 having the largest defect. The accumulation of dephosphorylated mutant PKC is consistent with increased PHLPP-mediated dephosphorylation of defectively autoinhibited PKC at the hydrophobic motif (*19*).

**Figure 4.**
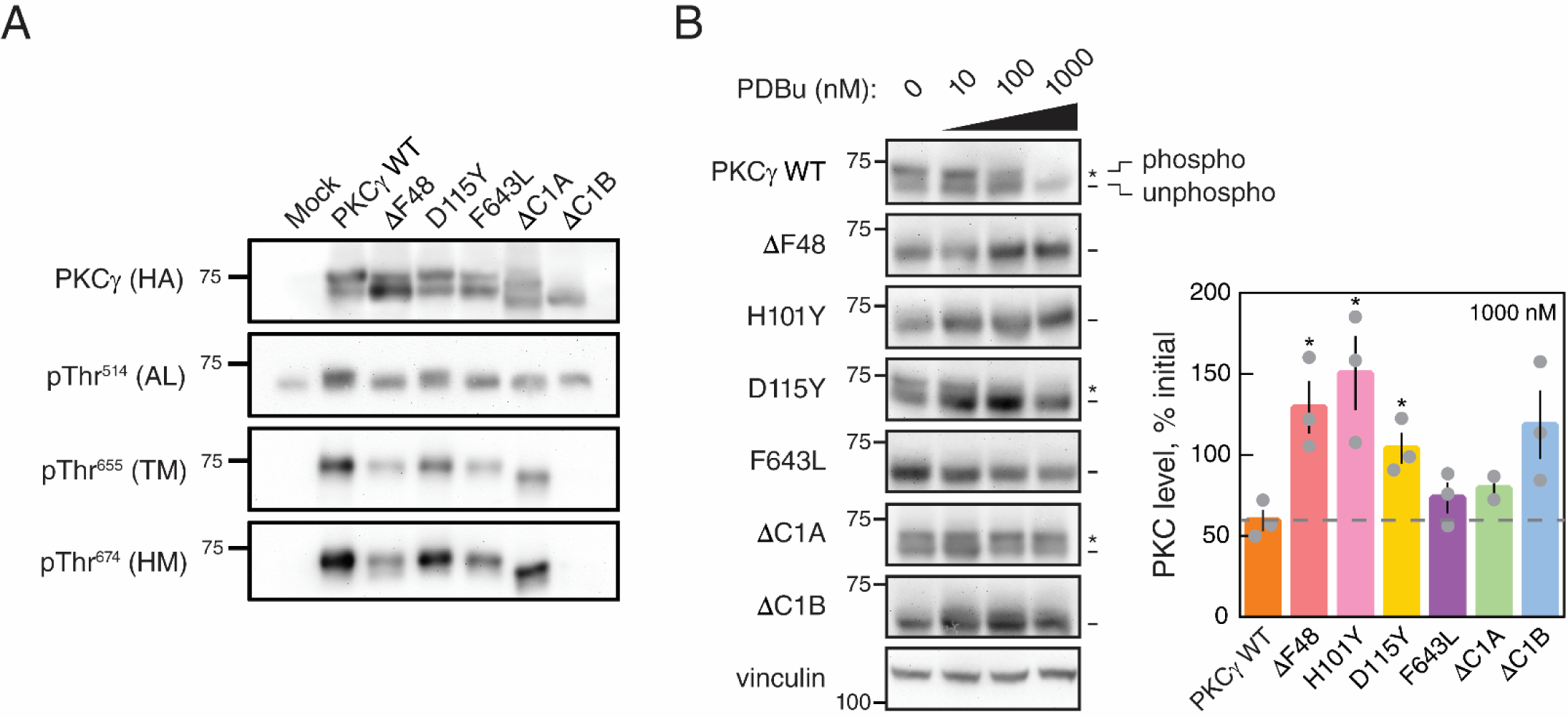
SCA14 mutants are resistant to phorbol ester-mediated downregulation. (A) Western blot of COS7 whole-cell lysates transfected with HA-tagged WT PKCγ, PKCγ lacking a C1A domain (ΔC1A), PKCγ lacking a C1B domain (ΔC1B), the indicated SCA14 mutants, or with empty vector (Mock). Membranes were probed with anti-HA (PKCγ) or phospho-specific antibodies. Data are representative of three independent experiments. (B) Western blot of lysates from COS7 cells transfected with HA-tagged WT PKCγ, PKCγ lacking a C1B domain (ΔC1B), PKCγ lacking a C1A domain (ΔC1A), or the indicated SCA14 mutants (left panel). COS7 cells were treated with the indicated concentrations of PDBu for 24 h prior to lysis. Endogenous expression of vinculin was also probed as a loading control. Data are representative of three independent experiments. *, phosphorylated species; -, unphosphorylated species. Quantification of total levels of PKC with 1000 nM PDBu (right panel) shown as a percentage of initial levels of PKC (0 nM) and represents mean ± S.E.M. Significance determined by Welch’s t-test (*P<0.05).

Given the increase in dephosphorylated species of SCA14 mutants, we next addressed whether these mutants were more susceptible to downregulation than WT PKCγ. COS7 cells overexpressing HA-tagged PKCγ WT, the indicated SCA14 mutants, ΔC1A, or ΔC1B were treated with increasing concentrations of PDBu for 24 hours (Fig. 4B) and PKC levels were probed by Western blot analysis of whole-cell lysates. Dephosphorylation of WT PKCγ was observed at the lowest concentration of PDBu (10 nM) as assessed by the accumulation of faster-mobility species, and this dephosphorylated species was degraded at the highest concentration of PDBu (1000 nM). Surprisingly, every C1 domain SCA14 mutant tested (ΔF48, H101Y, D115Y) was significantly more resistant to PDBu-mediated downregulation than WT PKCγ. The catalytic domain mutant F643L was also moderately less sensitive to PDBu downregulation than WT enzyme. Furthermore, ΔC1B, ΔF48, and H101Y levels increased with increasing concentrations of PDBu compared to levels in untreated cells. Although the C1B mutant D115Y was effectively dephosphorylated, the dephosphorylated species was resistant to degradation. In contrast, deletion of the C1A prevented dephosphorylation of the upper mobility, phosphorylated species, but allowed degradation of the faster mobility, dephosphorylated species. This demonstrates an uncoupling within the degradative pathway of PKC, such that a PKC that lacks a C1A domain is less susceptible to dephosphorylation, whereas a PKC without a functional C1B domain loses the ability to be degraded in a phorbol ester-dependent manner. Accumulation of mutant PKCγ in the Triton-insoluble fraction has previously been shown to be indicative of partially unfolded and degradation-resistant PKC (*40*). Probing for total PKC (HA) in either the Triton-soluble (Fig. S2A) or Triton-insoluble (Fig. S2B) fraction yielded a similar result, and revealed that the majority of the SCA14 mutants separate into the detergent-insoluble fraction following treatment of cells with 1000 nM PDBu. These results indicate that C1 domain mutants render PKC resistant to phorbol ester-mediated downregulation by impairing dephosphorylation (as observed upon deletion of C1A) or impairing degradation (as observed upon deletion of C1B). The kinase domain mutant F643L mirrored C1B domain mutations in resistance to phorbol ester-mediated degradation.

We next addressed whether SCA14-associated mutations altered the steady-state turnover of PKC in unstimulated cells. COS7 cells overexpressing HA-tagged PKCγ WT, the indicated SCA14 mutants, ΔC1A, or ΔC1B were treated with cycloheximide to prevent protein synthesis for increasing time and lysates were analyzed for PKC levels (Fig. 5A). PKCγ WT was remarkably stable, with a half-life of over 48 hours, as previously reported for other conventional PKC isozymes (*19*). In marked contrast, the ataxia mutants were considerably less stable, with half-lives of approximately 10 hours for mutations that had strong effects on autoinhibition (ΔF48, H101Y, F643L) and 20 hours for the D115Y mutation, which had a modest effect on autoinhibition (Fig. 5B). Deletion of the C1A or C1B domains (ΔC1A, ΔC1B) also had a strong effect on stability, consistent with decreased autoinhibition due to the loss of a regulatory domain. Thus, whereas SCA14 mutations render activated PKC resistant to phorbol ester-induced downregulation, they increase the steady-state turnover of unstimulated PKC.

**Figure 5.**
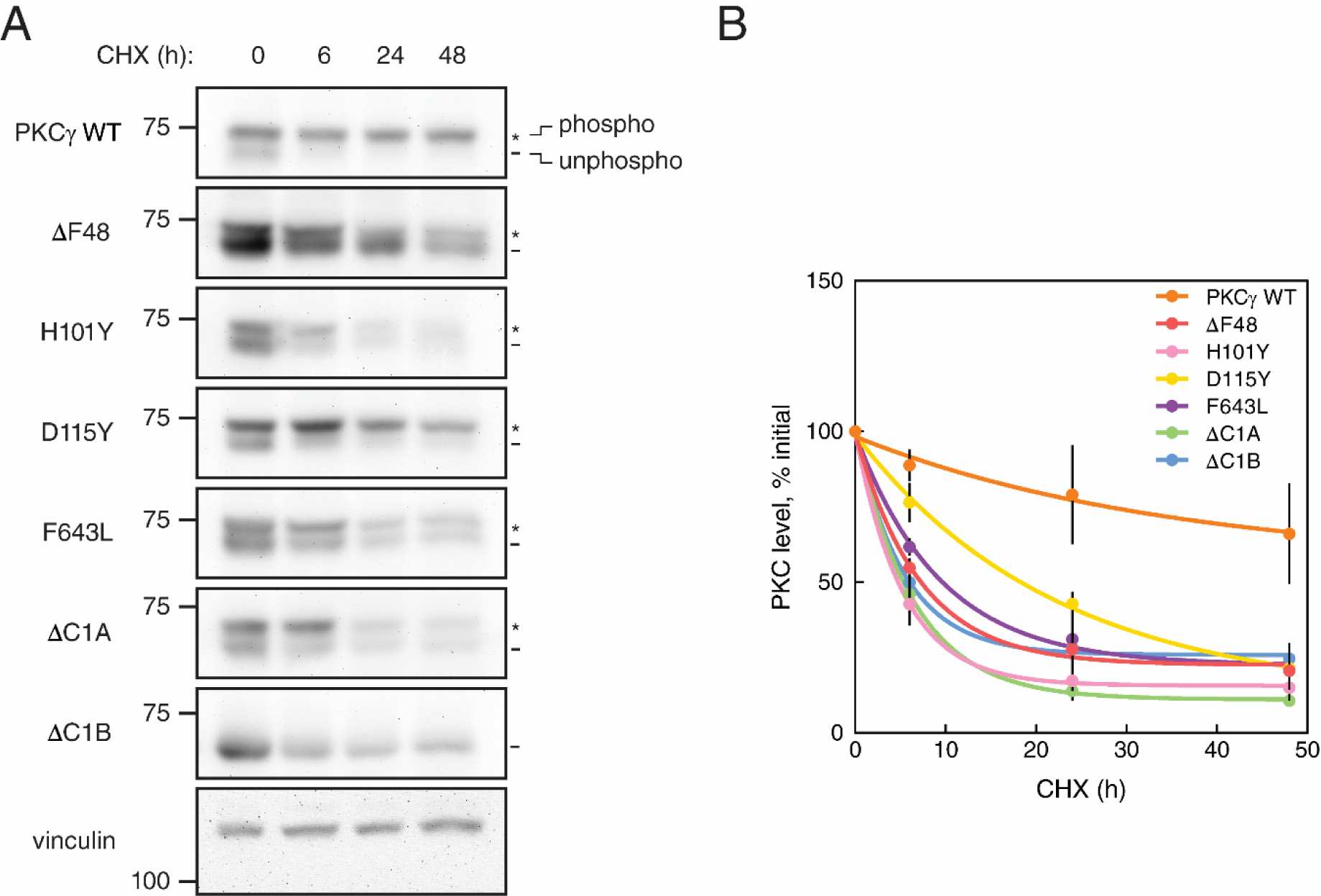
SCA14 mutants are more rapidly turned over in the presence of cycloheximide. (A) Western blot of lysates from COS7 cells transfected with HA-tagged WT PKCγ, PKCγ lacking a C1B domain (ΔC1B), PKCγ lacking a C1B domain (ΔC1A), or the indicated SCA14 mutants. COS7 cells were treated with 355 µM CHX for 0, 6, 24, or 48 hours prior to lysis. Endogenous expression of vinculin was also probed as a loading control. Data are representative of three independent experiments. *, phosphorylated species; -, unphosphorylated species. (B) Quantification of total levels of PKCγ at each time point shown as a percentage of initial levels of PKC (0 h) and represents mean ± S.E.M. Points were curve fit by non-linear regression.

### PKCγ C1A residue F48 is critical for proper autoinhibition and activation

The characterized SCA14 mutants displayed an increase in basal activity, and all but one retained the ability to have this elevated basal activity further enhanced in response to agonist stimulation (Fig. 2A-E). To gain insight into this uncoupling from agonist stimulated activity, we further characterized the deletion mutation in the C1A domain (ΔF48) whose activity was unresponsive to stimulation by UTP or PDBu, an uncoupling also observed upon deletion of the entire C1A domain (Fig. 2B). We first asked whether reducing the affinity of the pseudosubstrate for the active site pocket (Fig. 6A) or deleting the pseudosubstrate (Fig. 6B) would promote agonist- responsiveness of ΔF48. Mutation of arginine at the P-3 position to a glycine in WT (R21G) or ΔF48 (R21G ΔF48) PKCγ enhanced basal activity for both WT PKCγ and ΔF48 (Fig. 6C). However, UTP and PDBu caused additional activation of only the WT PKCγ with the pseudosubstrate mutation. While the pseudosubstrate mutation caused an even greater increase in basal activity of the SCA14 mutant, this still did not permit activation by UTP and PDBu (note that the small responses seen are those of the endogenous PKC). Similarly, deletion of the entire pseudosubstrate elevated basal activity even more for both WT and ΔF48, but further activation by PDBu was only observed for the PKCγ without the mutation in the C1A (Fig. 6D). Lastly, we addressed whether substitution (rather than deletion) of F48 restored agonist responsiveness. Mutation to either alanine (F48A) or the structurally more similar tyrosine (F48Y) restored autoinhibition to that observed for WT enzyme (Fig. 6E). However, while F48Y responded similarly to PDBu as WT PKCγ, F48A only partially rescued the WT response to PDBu. These data reveal that it is the loss of F48 that uncouples the pseudosubstrate from ligand engagement; substitution with Ala or Tyr may reduce activation kinetics and response to UTP, but still allows response to phorbol esters. We next examined a SCA14 deletion mutation at the corresponding position in the C1B domain (ΔF113) (Fig. 6F). Similar to ΔF48, ΔF113 had higher basal activity indicating impaired autoinhibition. However, the ΔF113 retained some responsiveness to phorbol esters, as evidenced by the increase in activity following PDBu stimulation. Thus, deletion of F48 in the C1A impairs autoinhibition but locks PKC in a conformation that prevents communication between the pseudosubstrate and membrane binding modules, whereas deletion of the corresponding F113 in the C1B impairs autoinhibition but allows more communication between the pseudosubstrate and membrane engagement.

**Figure 6.**
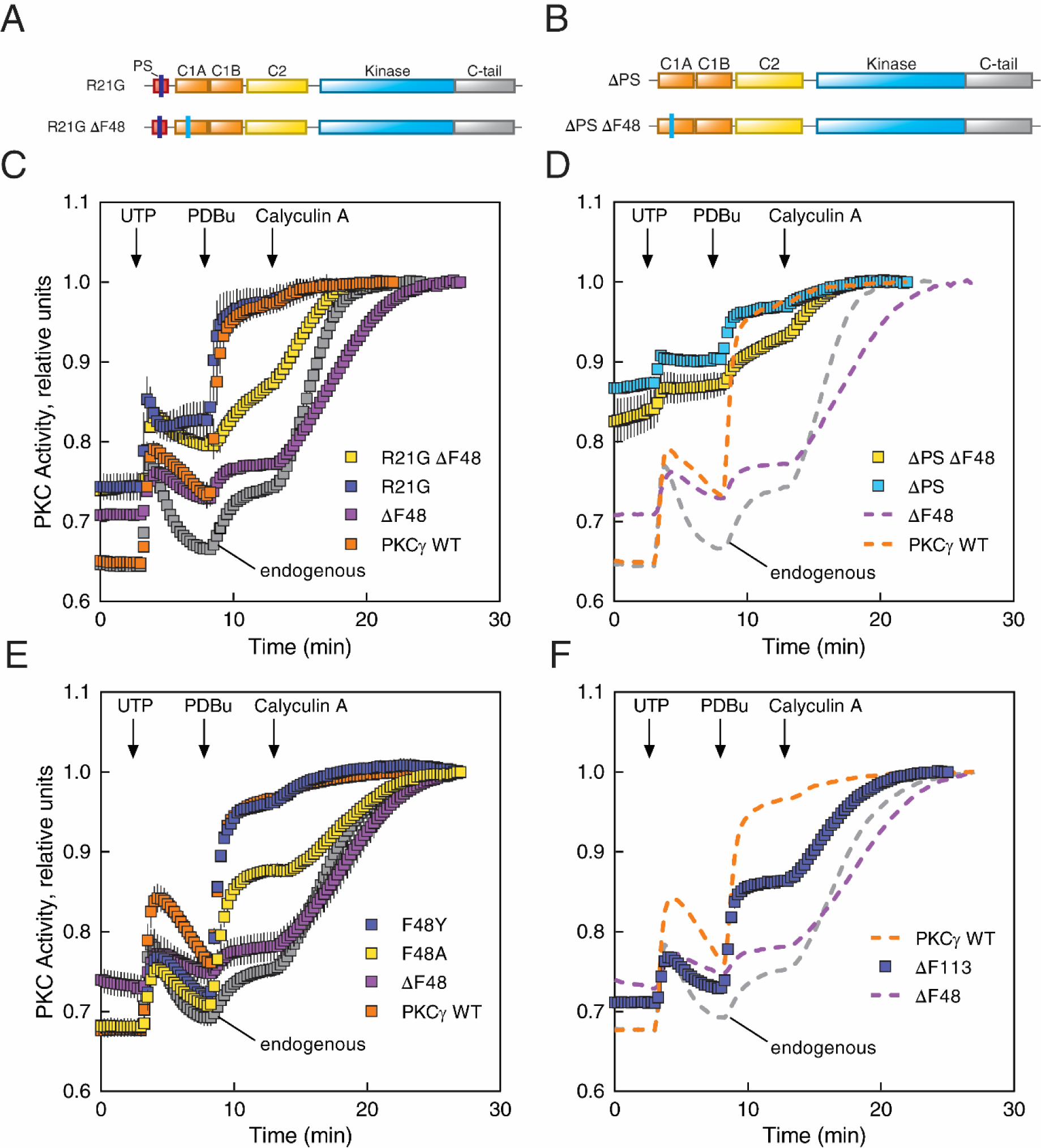
SCA14 mutant ΔF48 displays an abrogated response to agonists. (A) Domain structure of PKCγ constructs used in (**C**); mutated pseudosubstrate alone (R21G) or combined with F48 deleted (R21G ΔF48). (B) Domain structure of PKCγ constructs used in (**D**); deleted pseudosubstrate alone (ΔPS) or combined with F48 deleted (ΔPS ΔF48). (C) COS7 cells were transfected with CKAR2 alone (endogenous) or co-transfected with CKAR2 and the indicated mCherry-tagged PKCγ constructs in (**A**). PKC activity was monitored by measuring FRET/CFP ratio changes after addition of 100 µM UTP, 200 nM PDBu, and 50 nM Calyculin A. Data were normalized to the endpoint (1.0) and represent mean ± S.E.M. from at least two independent experiments (n ≥ 20 cells per condition). (D) COS7 cells were transfected with CKAR2 alone (endogenous) or co-transfected with CKAR2 and the indicated mCherry-tagged PKCγ constructs in (**B**). PKC activity was monitored by measuring FRET/CFP ratio changes after addition of 100 µM UTP, 200 nM PDBu, and 50 nM Calyculin A. Data were normalized to the endpoint (1.0) and represent mean ± S.E.M. from at least two independent experiments (n ≥ 20 cells per condition). PKCγ WT, ΔF48, and endogenous data are reproduced from (**C**) (dashed lines). (E) COS7 cells were transfected with CKAR2 alone (endogenous) or co-transfected with CKAR2 and the indicated mCherry-tagged PKCγ constructs. PKC activity was monitored by measuring FRET/CFP ratio changes after addition of 100 µM UTP, 200 nM PDBu, and 50 nM Calyculin A. Data were normalized to the endpoint (1.0) and represent mean ± S.E.M. from at least three independent experiments (n ≥ 49 cells per condition). (F) COS7 cells were transfected with CKAR2 alone (endogenous) or co-transfected with CKAR2 and the indicated mCherry-tagged SCA14 mutants. PKC activity was monitored by measuring FRET/CFP ratio changes after addition of 100 µM UTP, 200 nM PDBu, and 50 nM Calyculin A. Data were normalized to the endpoint (1.0) and represent mean ± S.E.M. from at least two independent experiments (n ≥ 31 cells per condition). PKCγ WT, ΔF48, and endogenous data are reproduced from (**E**) (dashed lines).

To validate whether the ΔF48 protein has lost the ability to be allosterically activated, we examined the activity of pure protein *in vitro* in the absence and presence of Ca^2+^ and lipid. GST-tagged PKCγ WT or ΔF48 produced in insect cells using a baculovirus expression system was purified to homogeneity (Fig. 7A) and activity was measured in the absence (non-activating conditions) or presence (activating conditions) of Ca^2+^ and multilamellar lipid structures (Fig. 7B). The activity of WT PKCγ was stimulated approximately 10-fold by Ca^2+^ and lipid, as reported previously (*41*), reflecting effective autoinhibition. In contrast, the specific activity of ΔF48 was approximately 3-fold higher than that of WT enzyme in the absence of cofactors, indicating impaired autoinhibition. Furthermore, addition of Ca^2+^/lipid had no effect on the activity of the ΔF48 mutant. Taken together with the activity data in live cells, these results establish that the ΔF48 C1A domain 1] has reduced autoinhibition, and 2] is locked in a conformation that prevents communication between the pseudosubstrate and the membrane-binding regulatory domains.

**Figure 7.**
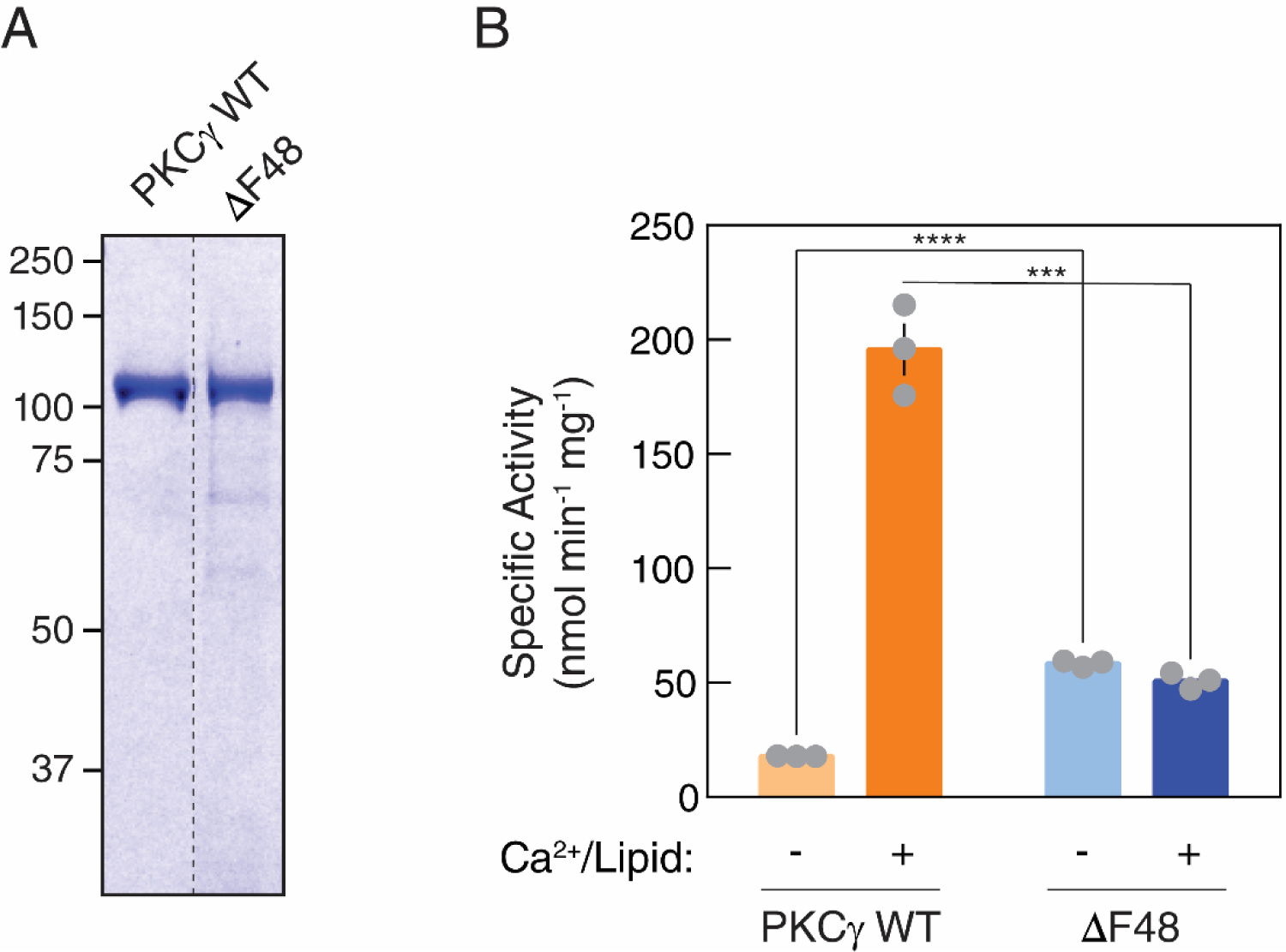
Purified ΔF48 exhibits increased activity compared to WT PKCγ under non- activating conditions. (A) Coomassie Blue-stained SDS-PAGE gel of purified GST-PKCγ WT or ΔF48. (B) *In vitro* kinase assays of purified GST-PKCγ WT or ΔF48 (6.1 nM per reaction). PKC activity was measured under non-activating conditions (EGTA, absence of Ca^2+^ or lipids) or activating conditions (presence of Ca^2+^ and lipids). Data are graphed in nanomoles phosphate per minute per milligram GST-PKC. Data represent mean ± S.E.M. from three independent experiments (n = 9 reactions per condition). Significance determined by multiple comparison *t*-tests (Holm-Sidak method) (***P<0.001, ****P<0.0001).

### Altered phosphoproteome in cerebellum of mice harboring SCA14-associated PKCγ mutation

The SCA14 C1 domain mutants tested all displayed increased basal activity (Fig. 2A) and resistance to phorbol ester downregulation in cell-based studies (Fig. 4B). To address whether this leaky activity altered the phosphoproteome in a physiological setting, we analyzed the cerebella from transgenic mice expressing human PKCγ WT or H101Y, or control C57BL/6 background mice via mass spectrometry (Fig. 8A). We quantified nearly 7000 unique proteins, from which 914 contained quantifiable phosphopeptide results across all samples. After correction for protein amount, the phosphoproteomics data identified a total of 195 phosphopeptides on 166 unique proteins demonstrating altered phosphorylation in the cerebella of H101Y-expressing mice compared to those expressing PKCγ WT, with 135 phosphopeptides significantly increasing in abundance and 60 phosphopeptides significantly decreasing in abundance in H101Y mice (Fig. 8B). Changes in phosphopeptide abundance were corrected by dividing phosphopeptide relative abundance by the corresponding protein abundance. Statistical significance was determined using a ranking method that simultaneously considers fold change and p-value (*42*) setting the α-value less than or greater than .05. Strikingly, 30 of the significantly decreased phosphopeptides were contained within neurofilament proteins (Fig. 8B, light blue circles), consistent with a general reduction in neurofilament phosphorylation in H101Y mouse cerebellum. Furthermore, the phosphorylation of one of the major kinases of neurofilaments glycogen synthase kinase 3 beta (GSK3β) (*43, 44*), was increased on an inhibitory site, Ser389 (Fig. 8C, left) (*45*). Our analyses also revealed an increase in phosphorylation at two sites (Ser22 and Ser26) on a single phosphopeptide of diacylglycerol kinase θ (DGKθ), which catalyzes the phosphorylation of DG to phosphatidic acid, in the H101Y mice, consistent with either direct or indirect regulation of DGKθ by PKCγ (Fig. 8C, right). Gene ontology analysis via DAVID GO (*46, 47*) (Fig. 8D) revealed that phosphopeptides with increased abundance in H101Y-expressing mice were primarily involved in processes related to axon extension, neural development, and cytoskeletal organization, and, similarly, phosphopeptides with decreased abundance in H101Y mice were mainly involved in neurofilament organization and axon development. This analysis suggests H101Y-expressing mice display dysregulation of signaling pathways involved in developing and maintaining neuron cytoskeletal structure and function, which may be regulated upstream by PKCγ.

**Figure 8.**
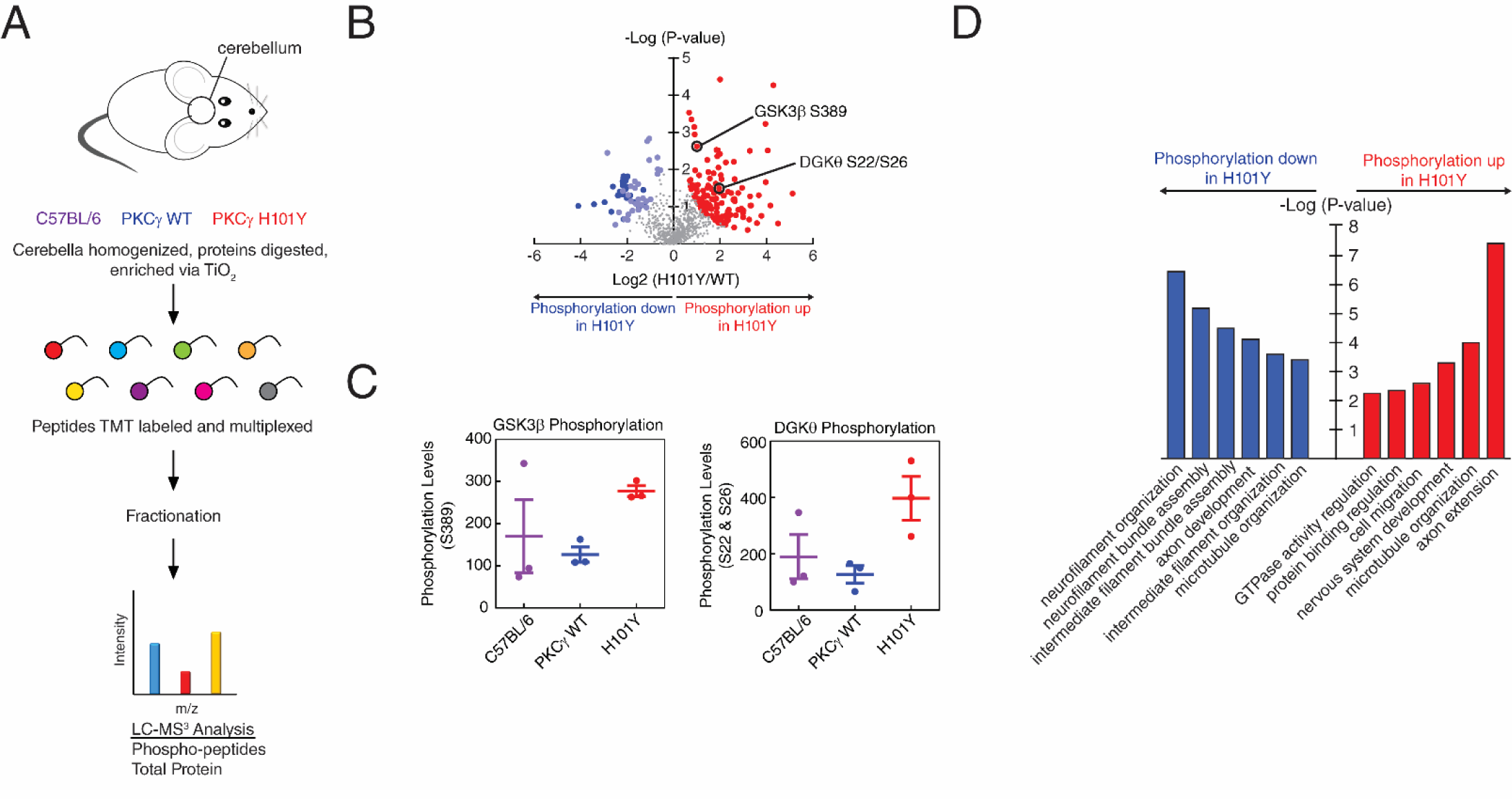
Phosphoproteomics analysis from cerebella of mice expressing human WT or H101Y PKCγ transgene. (A) Experimental design for processing of mouse tissue and proteins. Cerebella from B6 background (purple), PKCγ WT transgenic (blue), and PKCγ H101Y transgenic (red) mice were subjected to phosphoprotemics analysis. 6893 total proteins were quantified in the standard proteomics and 914 quantifiable phosphopeptides were detected in the phosphoproteomics. After correction for protein expression, 195 phosphopeptides on 166 unique proteins were identified in H101Y-expressing mice. (B) Volcano plot of phosphopeptide replicates of cerebella from WT and H101Y transgenic mice. Graph represents the log-transformed p-values (Student’s t-test) linked to individual phosphopeptides versus the log-transformed fold change in phosphopeptide abundance between WT and H101Y cerebella. Color represents phosphopeptides with significant changes in p-value and fold change; red, increased phosphorylation in H101Y mice; blue, lower phosphorylation in H101Y mice (dark blue indicates significantly decreased neurofilament phosphopeptides, light blue indicates all other significantly decreased phosphopeptides). (C) Graphs representing quantification of either a GSK3β phosphopeptide (left) or a DGKθ phosphopeptide (right) from the volcano plot in (**B**) in cerebella from C57BL/6 (purple), WT (blue), and H101Y (red) mice. (D) Gene ontology analysis of significantly increased (red) or decreased (blue) phosphopeptides representing significantly changed biological processes.

### Conventional PKC C1 domains are protected from mutation in cancer

We have previously shown that cancer-associated mutations in conventional PKC isozymes are generally loss-of-function (*48*), with mutations that impair autoinhibition triggering degradation by a PHLPP-mediated quality control mechanisms (*19*). However, SCA14 mutations, which occur with high frequency in the C1 domains, impair autoinhibition without triggering downregulation. None of the identified SCA14 mutations are currently annotated in cancer data bases such as cBioPortal (*49, 50*). Thus, we assessed whether the frequency of cancer-associated mutations in conventional PKC isozymes is lower in the C1 domains compared to the C2 domain. The number of missense mutations at each aligned residue position of PKCα, β, and γ was obtained from GDC Data Portal (Fig. 9A) (*51*) and the total mutation frequency within each domain (number of mutations per residues in the domain) was analyzed (Fig. 9B, left). The mutational frequency of the C1 domains was approximately half that of the C2 domain when all three conventional isozymes were analyzed together. Furthermore, we compared mutation frequencies of the C1B domain to all other domains and found that the C1B has significantly lower missense mutation frequency than other domains in PKC (Fig. 9B, right). Interestingly, analysis of the individual isozymes revealed that the C1A domain of PKCα was more protected from mutation than the C1B domain (Supplementary Table 1). Importantly, our analysis suggests that the C1B domain, a mutational hotspot in SCA14, is more protected overall from mutation in cancer compared to other domains.

**Figure 9.**
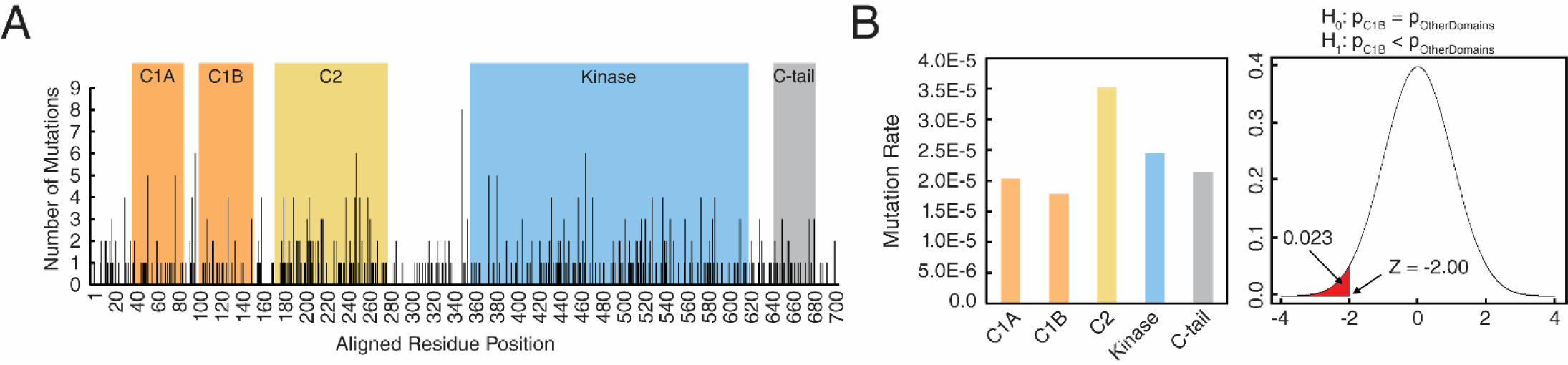
Statistical analysis of cancer mutation rate in PKC isozymes shows that C1B domain is protected from mutation. (A) Bar chart represents the number of cancer missense mutations (obtained from GDC Data Portal (*51*)) at each aligned residue position of conventional PKCs, including PKCα, β, and γ. Domains annotated by Pfam (*80*) are highlighted. (B) Bar chart shows the cancer mutation rate within each domain, which is defined by the total number of cancer missense mutations divided by the number of patients (10,189 patients) and the number of residues in the domain (left panel). A two-proportion z-test shows the cancer mutation rate of C1B is significantly lower than that of other domains (right panel), p-value = 0.023.

### Age of SCA14 onset inversely correlates with the degree of impaired PKCγ autoinhibition

To understand the degree to which the enhanced basal activity of the SCA14 mutants may contribute to disease, we plotted the level of biochemical defect (basal activity) against the age of onset of disease in the patients with the respective variants (*12, 52–57*) (Fig. 10A). This revealed that the degree of biochemical defect is directly proportional to disease severity: C1 domain mutants with high basal activity, such as V138E and ΔF48, were associated with an age of disease onset in early childhood (high disease severity), whereas those with lower levels of autoinhibitory defect, such as D115Y, were associated with an older age of onset (lower disease severity). The R^2^ value of approximately 0.76 supported an association between disease severity and degree of impaired autoinhibition.

**Figure 10.**
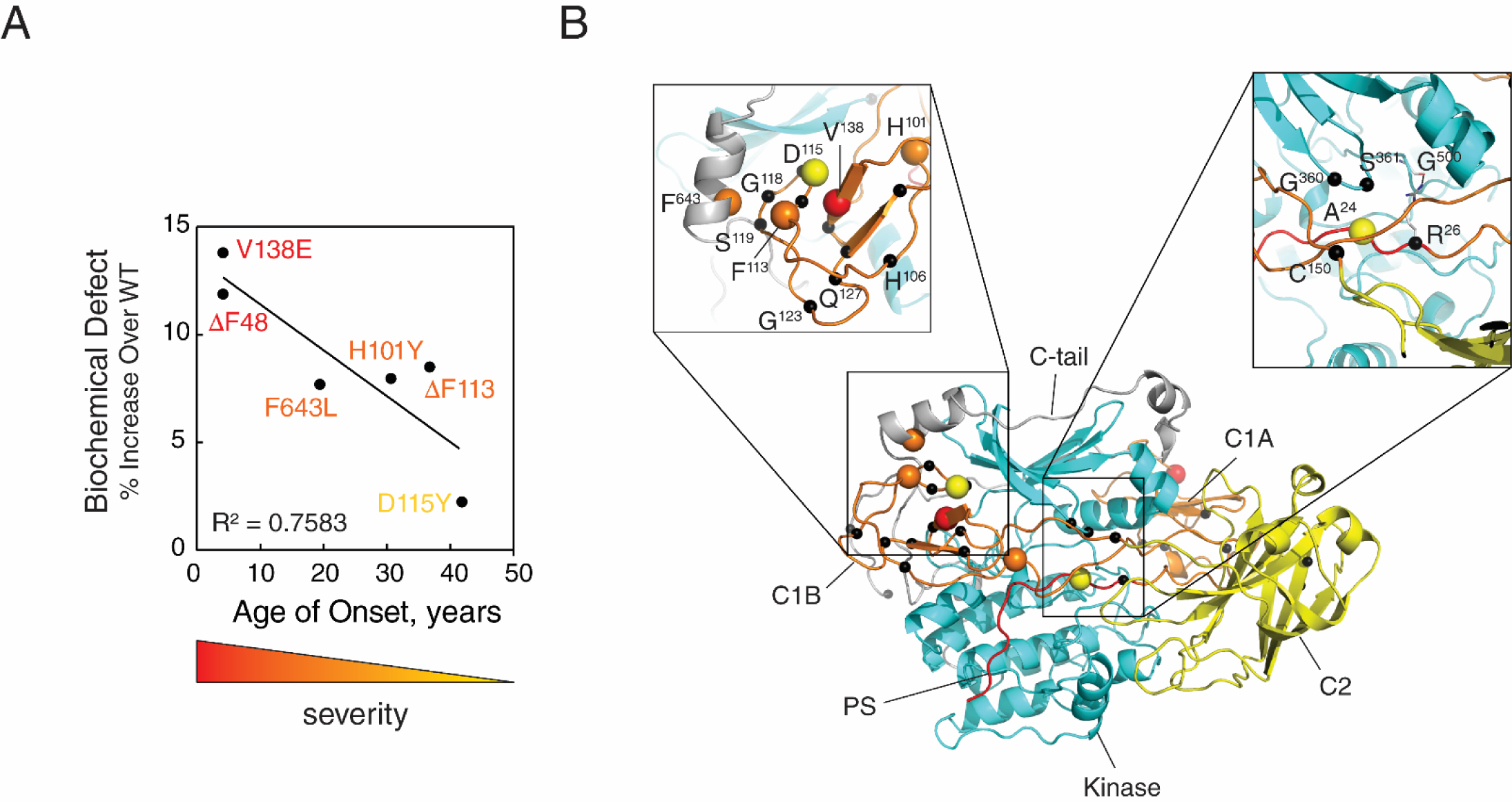
Degree of ataxia mutant biochemical defect correlates with SCA14 severity. (A) Graph of the indicated SCA14 mutant basal activities from Fig. 2B, C, E, and Fig. 6F plotted against age of disease onset in patients (*age of disease onset was reported as ‘early childhood’). Mutants are color-coded from red to yellow (most to least severe based on age of onset). (B) PKCγ model based on the previously published model of PKCβΙΙ (*16*). Indicated SCA14 mutations are represented as black spheres; the five mutations presenting in (A) are color coded by disease severity.

Lastly, we used a homology model for the architecture of conventional PKC isozymes (*16*) to predict where the 54 known SCA14-associated mutations (Fig. 1A) would occur within the 3- dimensional structure of PKCγ (Fig. 10B). In the autoinhibited conformation, the kinase adopts a compact conformation with the regulatory modules packed against the kinase domain and C-tail, and the pseudosubstrate segment (red) in the substrate binding cavity. Notably, many of the SCA14 mutations are predicted to exist either at an interface between the C1B and kinase domain (e.g. D115Y) or between the C1B domain and the C-tail (e.g., F643L). In particular, F643 is part of the conserved NFD motif, a key regulatory determinant of AGC kinases (*58*), which anchors the C1B in place (Fig. 10B, left inset) (*59*). Additionally, two mutations (A24T and R26G) are located in the pseudosubstrate, both of which are predicted to disrupt autoinhibition. The first, A24T, occurs at the phospho-acceptor site, which likely introduces a phosphorylation site, whereas R26G may disrupt a possible H-bond to G500 of the conserved DFG motif in the kinase domain (Fig. 10B, right inset). Only the two mutations in the C2 domain (I173S and H174P) were not at an interface with the kinase domain or regulatory domains. Thus, our model indicates that almost all SCA14 mutations target the C1 domains and their interfaces with the rest of the protein.

## Discussion

An abundance of germline variants in PKCγ are causal in SCA14, yet establishing whether a unifying mechanism accounts for the defect in these aberrant enzymes has remained elusive. Here we show that SCA14 PKCγ mutations in every domain of PKC (pseudosubstrate, C1A, C1B, C2, kinase, C-tail) display a shared autoinhibitory defect that leads to enhanced basal activity. Furthermore, by analyzing a mutant that uncouples pseudosubstrate regulation from phorbol ester binding, we show that increased basal signaling, rather than changes in agonist-evoked signaling, is the determinant associated with the ataxic phenotype. Remarkably, the degree of biochemical defect of the C1 domain mutants correlated inversely with age of onset of the disease. Thus, whereas previous studies have proposed a variety of mechanisms that may be involved in the cerebellar degeneration that is characteristic of ataxia (*23, 28–34*), our data showing a direct correlation between enhanced basal activity of PKCγ with disease severity are consistent with a model in which aberrant signaling by PKCγ in the absence of second messengers is the driver behind SCA14.

Disruption of autoinhibition of conventional PKC isozymes, either by mutation or by prolonged activation, as occurs with phorbol esters, results in unstable enzyme that is dephosphorylated and degraded (*60*). Indeed, this is a common mechanism for loss-of-function in cancer (*19*). Here, we show that mutations in the C1A or C1B domains, as well as deletion of either domain, renders PKCγ insensitive to phorbol ester-mediated downregulation. Thus, C1 mutations represent a susceptibility that allows for deregulated PKC activity without the paradoxical loss-of-function accompanying the ‘open’ conformation of PKC. This study also revealed that the two steps of downregulation can be uncoupled: the C1A domain is necessary for the first step in downregulation (dephosphorylation), and the C1B domain is necessary for the second step in downregulation (degradation). The finding that the C1 domain mutations evade downregulation provides an explanation for why these domains harbor the highest number of SCA14 mutations.

In this study, we identified a previously unknown SCA14-associated variant in the C1B domain of PKCγ (D115Y). Patients harboring this variant developed symptoms of the disease in their 40s, consistent with the mild biochemical defect observed in our study. Introduction of this mutation into the C1B domain of PKCδ has been shown not to significantly affect the affinity of the isolated domain for phorbol ester binding (*35*), however this residue is predicted to interface with the kinase domain (Fig. 10B). Thus, mutation to tyrosine could break interdomain interactions to favor the ‘open’ conformation because of the bulkier side chain of tyrosine compared to aspartate, and the loss of negative charge. Indeed, the phorbol ester-dependent translocation of D115Y PKCγ was considerably greater than that of WT PKCγ, consistent with a more exposed C1B domain. Thus, D115Y unmasks the C1B domain to modestly enhance basal signaling, resulting in a less severe pathology than C1B mutations that have more profound impairment on autoinhibitory constraints.

Although the C1 domain ataxia mutations conferred resistance to phorbol ester-mediated downregulation, the steady-state turnover of the mutants was enhanced compared to WT PKCγ. This uncoupling of agonist-dependent turnover and basal turnover has been reported previously. For example, the E3 ligase, RINCK was shown to promote PKC ubiquitination and degradation under non-activating conditions, however, phorbol ester-mediated downregulation was unaffected by siRNA knock-down of RINCK (*61*). Similarly, Leontieva and Black have identified two distinct pathways that mediate PKCα downregulation, one that is proteosome-dependent and one that is not (*62*). Taken with the results presented here, these data suggest that separate degradation pathways exist which affect passive turnover of basal PKC levels and degradation of activated PKC, respectively. How this increased basal turnover affects the steady-state levels of PKCγ in the disease awaits further studies.

A recurrent mutation in SCA14 is deletion of a Phe on the ligand binding loop of the C1 domains: ΔF48 in the C1A and ΔF113 in the C1B. Each mutation has the same effect on the autoinhibition of PKCγ as deletion of the entire domain, suggesting that deletion of this specific amino acid is functionally equivalent to loss of the domain. In the case of the C1A domain, this mutation (or deletion of the C1A) destroys communication between the pseudosubstrate and the C1B-C2 membrane-targeting modules. Thus, although the ΔF48 mutant is able to translocate to membranes when cells are treated with phorbol esters, this membrane engagement of the C1B domain does not allosterically activate PKC as it does for WT PKCγ. The ΔF48 mutation significantly impairs autoinhibition, and patients with this variant develop disease symptoms at a young age. This finding is strong evidence that enhanced basal signaling, and not an increase in agonist-evoked signaling, is the defect in SCA14. For the same mutation in the C1B domain, basal signaling is also enhanced, but communication with the pseudosubstrate is retained. As a result, phorbol ester stimulation further activates the enzyme, presumably by engagement of the C1A domain on membranes to release the pseudosubstrate. In summary, deletion of this conserved Phe in either the C1A or C1B inactivates the domain, with its loss in the C1A abolishing communication with the rest of the enzyme. Thus, the ΔF48 mutant is ‘frozen’ in a partially active conformation and cannot be allosterically activated, uncoupling it from DG and Ca^2+^ signaling.

What about mutations outside the C1 domains? Mutations that reduce the affinity of the pseudosubstrate for the active site destabilize PKC, promoting dephosphorylation and degradation (*19*). Yet four SCA14 mutations have been identified in the pseudosubstrate. Shimobayashi and Kapfhammer have provided key insight to this paradox by their analysis of a transgenic mouse harboring a mutation in the pseudosubstrate, A24E (*26*). This mutation, which caused an ataxic phenotype in mice and impaired Purkinje cell maturation, greatly reduced the stability of the enzyme and decreased steady-state levels approximately 10-fold compared with levels in WT mice. However, the unrestrained activity of the aberrant PKC that was present was sufficient to cause an increase in substrate phosphorylation in the cerebellum of these mice. Thus, although this PKC is unstable and steady-state levels are reduced, the basal activity is sufficiently elevated to drive the ataxic phenotype. Although our biochemical studies reveal that the pseudosubstrate mutations have the highest impaired autoinhibition of all the mutations studied (Fig. 2A), the age of onset for the disease in humans is relatively late (*54*). Taken together with the mouse model study, it is likely that the high basal activity is counterbalanced by the lower steady-state levels of the mutated enzyme to dampen the severity of the disease. This would be consistent with the model that enhanced basal activity is the driver of the phenotype. This enhanced basal activity likely produces neomorphic functions because the ‘leaky’ PKC activity is occurring in the cytosol and is not restricted to the plasma membrane.

Analysis of a SCA14 mouse model revealed that this ‘leaky’ PKC activity causes significant changes in the phosphorylation state of components of numerous processes in the cerebellum. Specifically, tandem mass tag (TMT) mass spectrometry-based proteomics utilizing phosphopeptide enrichment was used in the analysis of a transgenic mouse harboring the H101Y mutation revealed extensive alterations of the cerebellar phosphoproteome. The phosphorylation of a significant number of proteins increased and a significant fraction decreased. The decrease likely reflects the known regulation of phosphatase function by PKC (*63*). For example, the phosphorylation of PP2A B56δ at Ser566 was increased 2-fold in the H101Y cerebellum compared to WT; phosphorylation at this proposed PKC site has been reported to increase its activity (*64,65*). Most strikingly, the phosphorylation of both heavy, medium, and light neurofilament proteins decreased. Neurofilament proteins play key roles in growth of axons, with aberrations associated with neurodegeneration (*66*). One of the major kinases regulating neurofilaments is GSK3β (*43, 44*), which has been previously shown to be negatively regulated by PKC (*67*). Furthermore, our analysis revealed a two-fold reduction in the phosphorylation of GSK3β on an inhibitory site (S389) (*45*) in the cerebellum from the H101Y mice compared to WT. Underscoring neurofilaments as a target of aberrant PKCγ, gene ontology analysis identified neurofilament organization as the process with the most significant decrease in phosphopeptides. Conversely, of the peptides whose phosphorylation increased, axon development was the most significant class. Thus, leaky PKCγ activity causes significant rewiring of the cerebellar phosphoproteome.

The possibility that ‘leaky’ PKC signaling may be a driver in cerebellar ataxia, in general, is suggested by both unbiased network analyses and by specific mechanisms. First, a recent network analysis by Verbeek and colleagues identified alterations in synaptic transmission as one of the main shared mechanisms underlying genetically diverse SCAs (*68*). Our gene ontology analysis identified components that control synaptic transmission, such as axon extension and axon, as the most significant alterations resulting from aberrant PKCγ. Studies from Shirai and colleagues provide a potential mechanism: they recently reported that mice deficient in diacylglycerol kinase γ (DGKγ), which converts DG into phosphatidic acid and is regulated by PKC, display an ataxic phenotype (*69*). Additionally, defective dendritic development of Purkinje cells from these mice was reversed by inhibition of conventional PKC. Thus, elevation of basal DG levels drives Purkinje cell degeneration by a mechanism that depends on PKC activity. Purkinje cells from DGKγ knockout mice also display impaired induction of long-term depression (LTD), an important process that allows for cerebellar synaptic plasticity. Importantly, PKCα (*70*) but not PKCγ (*71*), is necessary for LTD in Purkinje cells, suggesting that the aberrant PKCγ is reducing PKCα function. Consistent with this possibility, Hirai and colleagues reported that LTD could not be induced in Purkinje cells expressing a SCA14 mutant of PKCγ (S119P) (*72*). Furthermore, co-expression of S119P PKCγ with PKCα resulted in decreased PKCα membrane residence time after depolarization-induced translocation. This led the authors to propose that increased (activating) phosphorylation of DGKγ by the SCA14 mutant of PKCγ would reduce DG levels, in turn reducing PKCα activity and impairing LTD induction. In support of a role for DGK in driving SCA14 pathogenesis, our phosphoproteomics analysis of cerebella from mice expressing human WT or H101Y PKCγ revealed that phosphorylation of DGKθ was significantly increased at Ser22 and Ser26 in the H101Y mice. Although the effect of these sites on DGKθ function has yet to be identified, PKC has been previously shown to positively regulate DGKθ translocation to plasma membrane (*73*). These data, taken with the results of previous studies, supports a model in which altered phosphorylation of multiple DGK isozymes may lead to DG depletion in Purkinje cells, thus reducing PKCα activation and membrane interaction. Although other mechanisms for such a dominant-negative effect on PKCα are also possible, one way in which enhanced basal activity in PKCγ may drive ataxia is by promoting DGK-dependent depletion of DG, ultimately impairing both PKCα activity and induction of LTD.

Cancer-associated mutations in conventional PKC isozymes are generally loss-of-function, whereas those identified in neurodegenerative disease are gain-of-function (*3, 4*). Given that the C1 domains provide a mechanism for gain-of-function without sensitivity to downregulation, we reasoned these domains may be under-mutated in cancer. Analysis of data from GDC Data Portal for all annotated mutations in conventional PKC isozymes (PKCα, PKCβ, PKCγ) revealed that the C1B domain exhibits a significantly lower mutation rate (1.79e-5 mutations per residue) than all other domains (Supplementary Table 1) (*51*). This was particularly strong for PKCβ and PKCγ, which exhibited a reduced mutation rate of 1.39e-5 mutations per residue. The finding that the C1B has a low mutational frequency in cancer is the converse of the enhanced mutational frequency in SCA14, supporting the model that gain of PKC function (rather than loss) is a driving force in neurodegenerative disease. We note that characterized germline mutations in PKCα in Alzheimer’s Disease do not impact autoinhibition, rather, they enhance the catalytic rate of the kinase domain such that a stronger response is evoked in response to agonist (*4, 6*). In contrast, SCA14 mutations increase basal activity, which drives the pathology. Thus, these two neurodegenerative diseases both have gain-of-function mutations in a conventional PKC, but Alzheimer’s Disease is associated with an enhancement in the acute, agonist-evoked activity of the enzyme, whereas SCA14 is driven by an enhancement in chronic, basal signaling. Strikingly, mutations that cause subtle increases in activity (either basal or agonist-evoked) are associated with both diseases, suggesting that small changes over a lifetime in long-lived cells such as neurons accumulate damage that eventually manifest in the disease, in the absence of other mutations.

To illuminate how SCA14 mutants might interfere with autoinhibition, we mapped mutations in the C1B and kinase domains on the PKCγ model, generated using the previously published model of PKCβII as a prototype (*16*). Indeed, many of these mutations exist in domains and at interfaces expected to interfere with autoinhibition of the kinase. However, this system is imperfect, as PKCγ architecture may be different from that of PKCβII, underscoring the need for a validated structure of PKCγ to accurately predict where and how SCA14 mutations affect PKCγ autoinhibition.

In summary, our study reveals that SCA14 mutations are uniformly associated with enhanced basal signaling of PKCγ, indicating that therapies that inhibit this enzyme may have therapeutic potential. In addition to identifying PKCγ as an actionable target in this neurodegenerative disease, our studies also provide a framework to predict disease severity in SCA14. Specifically, the direct correlation between the degree of impaired autoinhibition and disease severity allows prediction of patient prognosis of new mutations, such as the D115Y reported here. Lastly, given the direct regulation of PKCγ by intracellular Ca^2+^, and that many of the proteins mutated in other SCAs regulate Ca^2+^ homeostasis, one intriguing possibility is that enhanced PKCγ activity is not only central to SCA14 pathology, but is also at the epicenter of many other types of ataxia. This raises exciting possibilities for therapeutically targeting PKCγ in not just SCA14, but in many other subtypes of spinocerebellar ataxia.

## Materials and Methods

### Chemicals and antibodies

Uridine-5-triphosphate (UTP; cat #6701) and phorbol 12,13-dibutyrate (PDBu; cat #524390) were purchased from Calbiochem. Calyculin A (cat #9902) was purchased from Cell Signaling Technology. The anti-HA antibody (cat #11867423001) was purchased from Roche. The anti-phospho-PKCα/βII turn motif (pThr638/641; cat #9375) was from Cell Signaling Technology. Lipids used in kinase assays (DG, cat #800811C and PS, cat #840034C) were purchased from Avanti Polar Lipids. The anti-phospho-PKCγ hydrophobic motif (pThr674; cat# ab5797) antibody was from Abcam. The anti-phospho-PKCα/β/γ activation loop (pThr497/500/514) antibody was previously described (*44*). Ladder (cat #161-0394), bis/acrylamide solution (cat # 161-0156), and PVDF (cat# 162-0177) used for SDS-PAGE and Western blotting were purchased from Bio-Rad. Luminol (cat #A-8511) and p-coumaric acid (cat # C-9008) used to make chemiluminescence solution for western blotting were purchased from Sigma-Aldrich.

### Magnetic resonance imaging of ataxia patient brains

MRI Imaging was performed on a 1.5 Tesla Siemens MRI Scanner. Sagittal T1 Flair images were taken. Patients have consented to their anonymized scans being used in this publication.

### Plasmid constructs and mutagenesis

All mutants were generated using QuikChange site-directed mutagenesis (Agilent). PKC pseudosubstrate-deleted constructs were created by deletion of residues 19-36 via QuikChange mutagenesis (Agilent) using WT PKCγ, R21G, or ΔF48-containing mCherry-pcDNA3 plasmid. PKC C1A-, C1B-, and C2-deleted constructs were created by deletion of residues 36-75 (ΔC1A), 100-150 (ΔC1B), or 179-257 (ΔC2) via QuikChange mutagenesis (Agilent) using WT PKCγ mCherry- or HA-pcDNA3 plasmid. The C Kinase Activity Reporter 1 (CKAR1) (*27*) and C Kinase Activity Reporter 2 (CKAR2) (*26*) were previously described.

### Cell culture and transfection

COS7 cells were maintained in DMEM (Corning) containing 10% FBS (Atlanta Biologicals) and 1% penicillin/streptomycin (Gibco) at 37 °C in 5% CO2. Transient transfection was carried out using the Lipofectamine 3000 kit (ThermoFisher) per the manufacturer’s instructions, and constructs were allowed to express for 24 h for imaging, 24 h for CHX assays, or 48 h for PDBu downregulation assays and phosphorylation site Western blots.

### FRET imaging and analysis

COS7 cells were seeded into individual dishes, and imaging was performed under conditions and parameters previously described (*74*). Images were acquired on a Zeiss Axiovert microscope (Carl Zeiss Micro-Imaging, Inc.) using a MicroMax digital camera (Roper-Princeton Instruments) controlled by MetaFluor software (Universal Imaging Corp.). For CKAR activity assays, COS7 cells were co-transfected with the indicated mCherry-tagged PKCγ constructs and CKAR2 for 24 h before imaging, and cells were treated with 100 µM UTP, 200 nM PDBu, and 50 nM Calyculin A. For translocation assays, COS7 cells were co-transfected with the indicated YFP-tagged constructs and MyrPalm-mCFP (plasma-membrane targeted) (*38*) for 24 h before imaging, and cells were treated with 200 nM PDBu. For co-translocation assays, COS7 cells were co-transfected with mCherry-tagged PKCγ and YFP-tagged D115Y for 24 h before imaging, and cells were treated with 200 nM PDBu. Baseline images were acquired every 15 s for 3 min prior to treatment with agonists. For CKAR activity assays, all FRET ratios were normalized to the endpoint of the assay. Translocation assays are normalized to the starting point of the assay.

### Phorbol ester downregulation assay

COS7 cells were seeded in 6-well plates at 1.5x10^5^ cells per well. After 24 h, cells were transfected with indicated HA-tagged PKCγ constructs (100ng DNA per well) for 48 h before PDBu treatment. Cells were treated with 10 – 1000 nM PDBu or DMSO for 24 h. Cells were then washed with DPBS (Corning) and lysed in PPHB containing 50 mM NaPO4 (pH 7.5), 1% Triton X-100, 20 mM NaF, 1 mM Na4P2O7, 100 mM NaCl, 2 mM EDTA, 2 mM EGTA, 1 mM Na3VO4, 1 mM PMSF, 40 µg/mL leupeptin, 1 mM DTT, and 1 µM microcystin. For PDBu downregulation assays, whole-cell lysate was loaded on gels. For fractionation assays, Triton-insoluble pellets were separated from soluble fractions by centrifugation at 4 °C, then pellets were resuspended in buffer containing 25 mM HEPES (pH 7.4), 3mL 0.3M NaCl, 1.5 mM MgCl2, 1 mM Na3VO4, 1 mM PMSF, 40 µg/mL leupeptin, 1 mM DTT, and 1 µM microcystin. Benzonase was added to whole-cell lysates and Triton-insoluble fractions at 1:100 to digest nucleotides.

### Cycloheximide assay

COS7 cells were seeded in 6-well plates at 1.5x10^5^ cells per well. After 24 h, cells were transfected with indicated HA-tagged PKCγ constructs (100ng DNA per well) for 24 h before CHX treatment. Cells were treated with 355 µM or DMSO for 0, 6, 24, or 48 h. Cells were then washed with DPBS (Corning) and lysed in PPHB containing 50 mM NaPO4 (pH 7.5), 1% Triton X-100, 20 mM NaF, 1 mM Na4P2O7, 100 mM NaCl, 2 mM EDTA, 2 mM EGTA, 1 mM Na3VO4, 1 mM PMSF, 40 µg/mL leupeptin, 1 mM DTT, and 1 µM microcystin. Whole-cell lysate was loaded on gels. Benzonase was added to whole-cell lysates at 1:100 to digest nucelotides.

### Western blots

All cell lysates were analyzed by SDS-PAGE overnight on 6.5% big gels at 9 mA per gel to observe phosphorylation mobility shift. Gels were transferred to PVDF membrane (Bio-Rad) by a wet transfer method at 4 °C for 2 h at 80 V. Membranes were blocked in 5% BSA in PBST for 1 h at room temperature, then incubated in primary antibodies overnight at 4 °C. Membranes were washed for 5 min three times in PBST, incubated in appropriate secondary antibodies for 1 h at room temperature, washed for 5 min three times in PBST, then imaged via chemiluminescence (100 mM Tris pH 8.5, 1.25 mM Luminol, 198 μM coumaric acid, 1% H2O2) on a FluorChem Q imaging system (ProteinSimple). In western blots, the asterisk (*) indicates phosphorylated PKC species, while a dash (-) indicates unphosphorylated species.

### Purification of GST-PKC from Sf9 insect cells

Wild-type PKCγ and ΔF48 were cloned into the pFastBac vector (Invitrogen) containing an N-terminal GST tag. Using the Bac-to-Bac Baculovirus Expression System (Invitrogen), the pFastBac plasmids were transformed into DH10Bac cells, and the resulting bacmid DNA was transfected into Sf9 insect cells via CellFECTIN (ThermoFisher). Sf9 cells were grown in Sf-900 II SFM media (Gibco) in shaking cultures at 27 °C. The recombinant baculoviruses were harvested and amplified three times. Sf9 cells were seeded in 125mL spinner flasks at 1x10^6^ cells per mL and infected with baculovirus. After 2 days of infection, Sf9 cells were lysed in buffer containing 50 mM HEPES (pH 7.5), 1 mM EDTA, 100 mM NaCl, 1% Triton X-100, 100 μM PMSF, 1 mM DTT, 2 mM benzamidine, 50 μg/ml leupeptin, and 1 µM microcystin. Soluble lysates were incubated with GST-Bind resin (EMD Millipore) for 30 min at 4 °C, washed three times, then GST-PKCγ was eluted off the beads with buffer containing 50 mM HEPES (pH 7.5), 1 mM EDTA, 100 mM NaCl, 1 mM DTT, and 10 mM glutathione. Purified protein was concentrated with Amicon Ultra-0.5 mL centrifugal filters (50kDa cutoff; EMD Millipore) to 100 µL. Protein purity and concentration was determined using BSA standards on an 8% SDS-PAGE mini-gel stained with Coomassie Brilliant Blue stain.

### *In vitro* kinase activity assays

The activity of purified GST-PKCγ (6.1 nM) upon a MARCKS peptide substrate (Ac-FKKSFKL- NH2) was assayed as previously described (*75*). Reactions contained 20 mM HEPES (pH 7.4), 0.06 mg/mL BSA, 1.2 mM DTT, 100 µM ATP, 100 µM peptide substrate, and 5 mM MgCl2. For activating conditions, Ca^2+^ (final concentration of 100 µM) and multilamellar lipids containing 15 mol % PS and 5 mol % DG were added to the reaction mixes. For non-activating conditions, 1 M HEPES (pH 7.4) and 500 µM EGTA were added in volumes equal to those of the lipids and Ca^2+^ in activating reaction conditions. Upon addition of ^32^P-ATP (Perkin Elmer), reactions were allowed to proceed at 30 °C for 10 min.

### Mouse model and harvest of cerebella

Under an approved University of Washington Institutional Animal Care and Use Committee (IACUC) protocol, SCA14 mutant (H101Y) and wild type (WT) PKCγ transgenic (Tg) mice were generated using modified human-BAC constructs, where expression of human PKCγ is regulated by the endogenous human PRKCG promoter. Flanking neuronal-expressed genes were removed and an eGFP-tag was introduced to enable detection and visualization of the transgene. Purified PRKCG fragments were microinjected into C3H/C57BL6 hybrid oocyte pronuclei. RNA, western blot and immunohistochemical analyses all confirmed expression of the transgene. We interbred the lines to homozygosity for the transgenes (WT- and H101Y-PRKCG++). The level of expression of the transgene was lower than that of the endogenous PRKCG as detected by eGFP fluorescence, but by 3 months of age, H101Y-Tg++ mice demonstrated impaired rotarod performance compared to WT-PRKCG Tg mice. By 3 months of age, H101Y Tg++ lines demonstrated impaired rotarod performance compared to WT-PRKCG Tg mice. At 3-months of age, three mice of homozygous-Tg genotype and three C56BL/6 mice were sacrificed by cervical dislocation and cerebella were dissected, snap-frozen in liquid nitrogen and kept at -80°C until shipment on dry ice to UCSD for protein extraction and proteomics analysis.

### Mass spectrometry-based proteomics

Sample processing, phosphopeptide enrichment and mass spectrometry analysis followed methods described previously (*76*), but are described here briefly to highlight modifications. Snap frozen cerebellum (approximately 45 mg) were homogenized by bead beating at 37 degrees in lysis buffer (1 mL) composed of 3% SDS, 75 mM NaCl, 1 mM NaF, 1 mM β-glycerophosphate, 1 mM sodium orthovanadate, 10 mM sodium pyrophosphate, 1 mM PMSF and 1X Roche Complete mini EDTA free protease inhibitors in 50 mM HEPES, pH 8.5. Rough homogenates were then further subjected to probe sonication (Q500 QSonica sonicator with 1.6 mm microtip). To the protein mixture was added an equal volume of Urea (8 M in 50 mM HEPES). Samples were reduced and alkylated using dithiothreitol (5 mM) and iodoacetamide (15 mM) respectively. Proteins were precipitated using chloroform/methanol, dried, and resuspended in 1 M urea in 50 mM HEPES (600 µL). Proteins were then digested using LysC, followed by trypsin before purification by SepPak cartridges. Protein aliquots (50 µg) from each sample were lyophilized and stored at -80 ℃ for labeling and proteomic analysis, along with 7 µg per sample pooled to generate two bridge channels. From each sample 1 mg peptide was subjected to phospho-enrichment using 6 mg Titanium dioxide beads. Enriched peptides were desalted using solid-phase extraction columns, lyophilized and stored at -80 ℃ until labeling.

For both the phospho-enriched peptides and reserved peptides for proteomics, peptides were labeled with tandem mass tag (TMT) reagents, reserving the 126 and 131 mass labels for the two bridge channels. Labeled samples were then pooled into multiplex experiments and de-salted by solid phase extraction. Sample fractionation was performed using spin columns to generate eight fractions per multiplex experiment. Fractions were lyophilized, re-suspended in 5% formic acid/5% acetonitrile for LC-MS2/MS3 identification and quantification. LC-MS2/MS3 analysis were performed on an Orbitrap Fusion mass spectrometer and data processing was carried out using the ProteomeDiscoverer 2.1.0.81 software package as described previously (*76*).

### PKCγ model

The PKCγ model was built in UCSF Chimera 1.13.1 (*77*) with integrated Modeller 9.21 (*78*). The kinase domain was modelled using the structure of PKCβΙΙ (PDB: 2I0E) as a template. The structure of the C1B domain was modelled using the structure of the C1A of PKCγ (PDB: 2E73). The C1 domains were docked to the kinase domain according to the previously published model of PKCβ (*59*). The structure of the PKCγ C2 domain (PDB: 2UZP) was docked to the kinase domain and C1 domains complex using the PKCβΙΙ model as a starting point using ClusPro web server (*79*).

### Quantification and data analysis

FRET ratios for CKAR assays were acquired with MetaFluor software (Molecular Devices). Ratios were normalized to starting point or endpoint (1.0) as indicated in figure legends. Western blots were quantified by densitometry using the AlphaView software (ProteinSimple). Gene ontology was performed by DAVID GO (*46, 47*) and was background adjusted using the *Mus musculus* species background. Statistical significance was determined via unequal variances (Welch’s) *t*-test or multiple *t*-tests (with the Holm-Sidak method of determining significance) performed in GraphPad Prism 8 (GraphPad Software).

## Supplementary Materials

Fig. S1. CKAR2 has a larger dynamic range than CKAR1.

Fig. S2. SCA14 mutants resist phorbol ester-mediated downregulation in both the Triton soluble and insoluble fractions.

Supplementary Table 1. Cancer missense mutation frequency differs in each domain between conventional PKC isozymes.

## Supporting information

Phosphoproteomics Raw Data

## Acknowledgments

We thank Dr. Natasha Fullerton for help with image analysis and providing the image control of the MRI scans. We thank the laboratory of J. Zhang (UCSD) for generously providing the CKAR2. We thank Dr. James T. Yurkovich for advice on initial analysis of frequency of cancer-associated mutations. We also thank all members of our laboratories for their technical work and helpful comments on this manuscript.

## Author contributions

C.A.P. and M.T.K. performed the experiments. A.K. and S.S.T. performed the molecular modeling. C.L. and G.G. identified D115Y SCA14 mutation and performed and analyzed the magnetic resonance imaging of patients. D.H.C. and W.H.R. generated and phenotyped the mouse models of SCA14. M.M. and D.J.G. performed and analyzed the phosphoproteomics. L.H. and N.K. performed bioinformatics and statistical analyses of domain mutation rates in cancer. T.B. performed pilot experiments. C.A.P. and A.C.N. designed the experiments and wrote the manuscript and all authors edited the draft.

## Supplementary Materials

**Figure S1.**
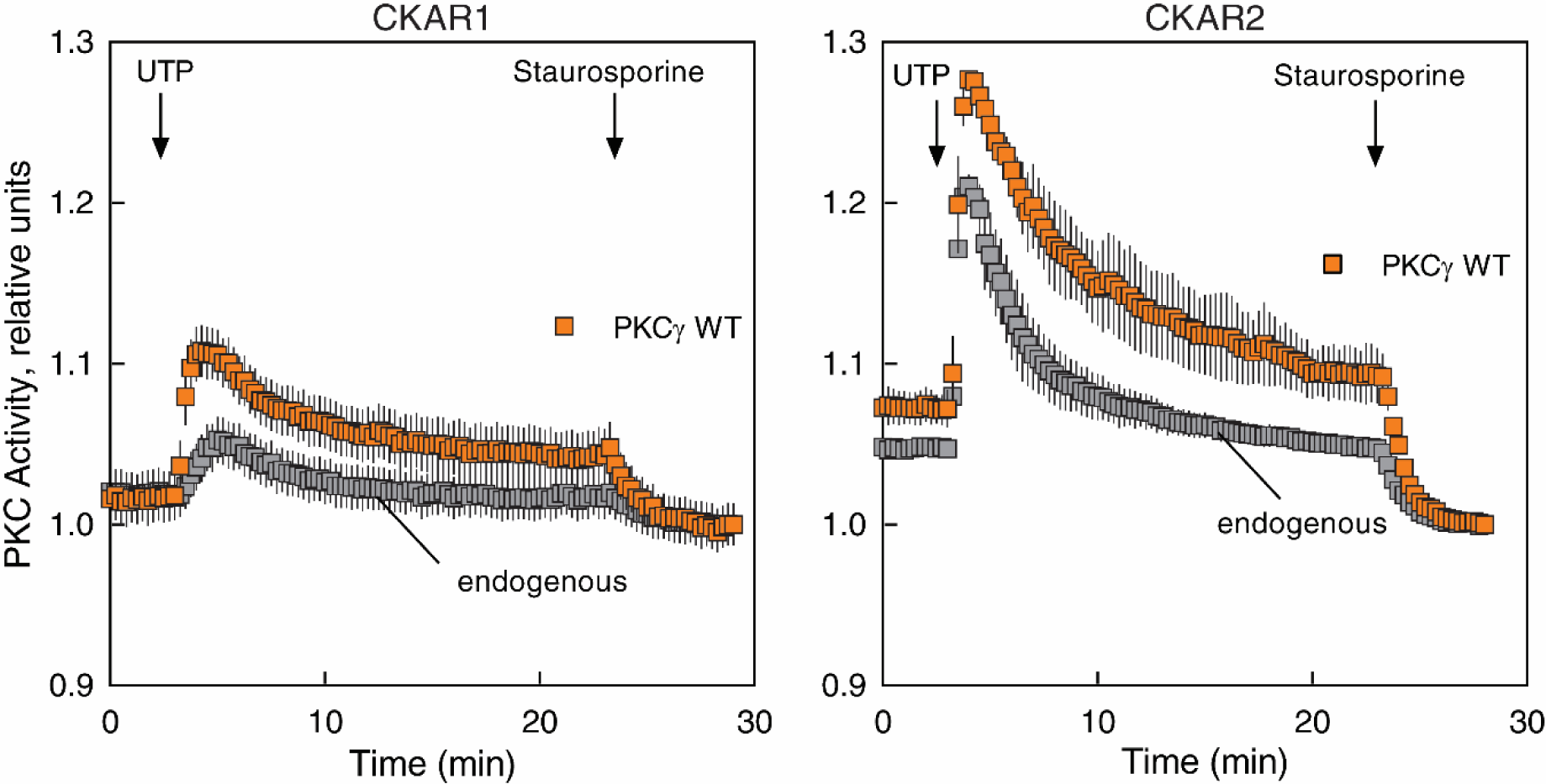
CKAR2 has a larger dynamic range than CKAR1. Left panel: COS7 cells were transfected with CKAR1 alone or co-transfected with CKAR1 and mCherry-PKCγ WT. PKC activity was monitored by measuring CFP/FRET ratio changes after addition of 100 µM UTP and 1 µM staurosporine. Data were normalized to the endpoint (1.0) and represent mean ± S.E.M. (n ≥ 15 cells per condition). Right panel: COS7 cells were transfected with CKAR2 alone or co-transfected with CKAR2 and mCherry-PKCγ WT. PKC activity was monitored by measuring FRET/CFP ratio changes after addition of 100 µM UTP and 1 µM staurosporine. Data were normalized to the endpoint (1.0) and represent mean ± S.E.M. (n ≥ 18 cells per condition).

**Figure S2.**
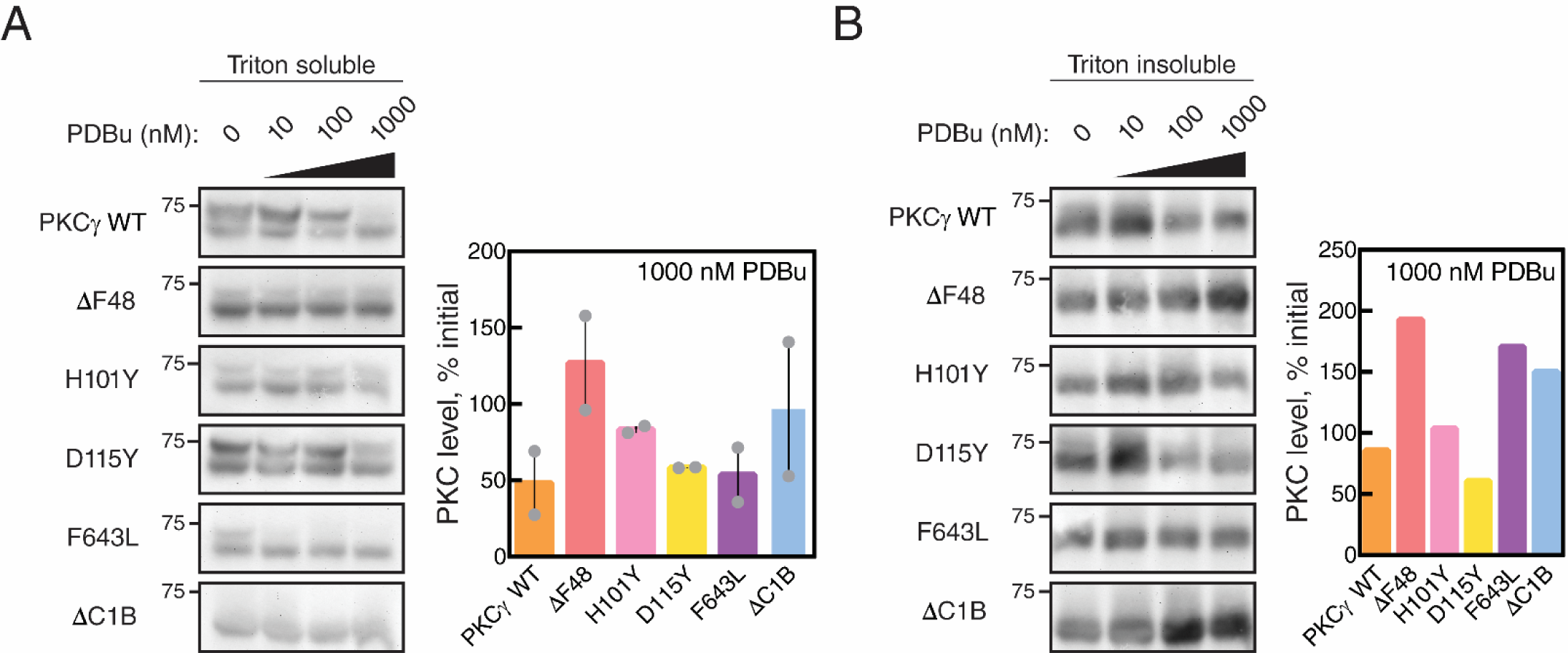
SCA14 mutants resist phorbol ester-mediated downregulation in both the Triton soluble and insoluble fractions. (A) Western blot (left) of Triton soluble lysate fractions from COS7 cells transfected with HA- tagged WT PKCγ, PKCγ lacking a C1B domain (ΔC1B), or the indicated SCA14 mutants. COS7 cells were treated with the indicated concentrations of PDBu for 24 h prior to lysis. Data is representative of two independent experiments. Quantification of total levels of PKC with 1000 nM PDBu (right) shown as a percentage of initial levels of PKC (0 nM) and represents mean ± S.E.M. (B) Western blot (left) of Triton insoluble lysate fractions from COS7 cells transfected with HA- tagged WT PKCγ, PKCγ lacking a C1B domain (ΔC1B), or the indicated SCA14 mutants. COS7 cells were treated with the indicated concentrations of PDBu for 24 h prior to lysis. Data represents one independent experiment. Quantification of total levels of PKC with 1000 nM PDBu (right) shown as a percentage of initial levels of PKC (0 nM).

**Supplementary Table 1.** Cancer missense mutation frequency differs in each domain between conventional PKC isozymes. Table represents the number of cancer missense mutations (obtained from GDC Data Portal (51)) in each domain of individual conventional PKC isozymes, including PKCα, β, and γ. * Total number of residues are calculated by the total domain length times 10,189 patients.

| Domain | Number of missense mutations |  |  |  | Total Domain Length | Total No. of Residues* | MT/Residues |
| --- | --- | --- | --- | --- | --- | --- | --- |
| | PKCα | PKCβ | PKCγ | Total | | | |
| C1A | 4 | 17 | 12 | 33 | 159 | 1,620,051 | 2.04e-5 |
| C1B | 14 | 9 | 6 | 29 | 159 | 1,620,051 | **1.79e-5** |
| C2 | 22 | 44 | 49 | 115 | 321 | 3,270,669 | 3.52e-5 |
| Kinase | 34 | 93 | 68 | 195 | 781 | 7,957,609 | 2.45e-5 |
| C-tail | 7 | 14 | 4 | 25 | 114 | 1,161,546 | 2.15e-5 |
| **Total** | **81** | **177** | **139** | **397** | **1,534** | **15,629,926** | **2.54e-5** |

## References and Notes

1. J. A. Callender, A. C. Newton, Conventional protein kinase C in the brain: 40 years later. Neuronal Signaling. 1, 20160005 (2017).

2. Y. Nishizuka, Protein kinase C and lipid signaling for sustained cellular responses. The FASEB Journal. 9, 484–496 (1995).

3. A. C. Newton, J. Brognard, Reversing the Paradigm: Protein Kinase C as a Tumor Suppressor. Trends in Pharmacological Sciences. 38 (2017), pp. 438–447.

4. J. A. Callender, Y. Yang, G. Lordén, N. L. Stephenson, A. C. Jones, J. Brognard, A. C. Newton, Protein kinase Cα gain-of-function variant in Alzheimer’s disease displays enhanced catalysis by a mechanism that evades down-regulation. Proceedings of the National Academy of Sciences of the United States of America. 115, E5497–E5505 (2018).

5. H. Tovell, A. C. Newton, PHLPPing the balance: restoration of protein kinase C in cancer. Biochemical Journal. 478 (2021), pp. 341–355.

6. S. I. Alfonso, J. A. Callender, B. Hooli, C. E. Antal, K. Mullin, M. A. Sherman, S. E. Lesné, M. Leitges, A. C. Newton, R. E. Tanzi, R. Malinow, Gain-of-function mutations in protein kinase Cα (PKCα) may promote synaptic defects in Alzheimer’s disease. Science Signaling. 9 (2016), doi:10.1126/scisignal.aaf6209.

7. Y. M. Sun, C. Lu, Z. Y. Wu, Spinocerebellar ataxia: relationship between phenotype and genotype – a review. Clinical Genetics. 90, 305–314 (2016).

8. M. Tada, M. Nishizawa, O. Onodera, Roles of inositol 1,4,5-trisphosphate receptors in spinocerebellar ataxias. Neurochemistry International. 94, 1–8 (2016).

9. J. Liu, T. S. Tang, H. Tu, O. Nelson, E. Herndon, D. P. Huynh, S. M. Pulst, I. Bezprozvanny, Deranged calcium signaling and neurodegeneration in spinocerebellar ataxia type 2. Journal of Neuroscience. 29, 9148–9162 (2009).

10. B. L. Fogel, S. M. Hanson, E. B. E. Becker, Do mutations in the murine ataxia gene TRPC3 cause cerebellar ataxia in humans? Movement Disorders. 30 (2015), pp. 284–286.

11. L. M. Watson, E. Bamber, R. P. Schnekenberg, J. Williams, C. Bettencourt, J. Lickiss, K. Fawcett, S. Clokie, Y. Wallis, P. Clouston, D. Sims, H. Houlden, E. B. E. Becker, A. H. Németh, Dominant Mutations in GRM1 Cause Spinocerebellar Ataxia Type 44. American Journal of Human Genetics. 101, 451–458 (2017).

12. D. H. Chen, Z. Brkanac, C. L. M. J. Verlinde, X. J. Tan, L. Bylenok, D. Nochlin, M. Matsushita, H. Lipe, J. Wolff, M. Fernandez, P. J. Cimino, T. D. Bird, W. H. Raskind, Missense mutations in the regulatory domain of PKCγ: A new mechanism for dominant nonepisodic cerebellar ataxia. American Journal of Human Genetics. 72, 839–849 (2003).

13. N. Saito, Y. Shirai, Protein kinase Cγ (PKCγ): Function of neuron specific isotype. Journal of Biochemistry. 132, 683–687 (2002).

14. F. Metzger, J. P. Kapfhammer, Protein-kinase C: Its role in activity-dependent Purkinje cell dendritic development and plasticity. Cerebellum. 2 (2003), pp. 206–214.

15. A. C. Newton, Protein kinase C: perfectly balanced. Critical Reviews in Biochemistry and Molecular Biology. 53, 208–230 (2018).

16. A. C. Jones, S. S. Taylor, A. C. Newton, A. P. Kornev, Hypothesis: Unifying model of domain architecture for conventional and novel protein kinase C isozymes. IUBMB Life. 72, 2584–2590 (2020).

17. J. H. Evans, D. Murray, C. C. Leslie, J. J. Falke, Specific translocation of protein kinase Ca to the plasma membrane requires both Ca2+ and PIP2 recognition by its C2 domain. Molecular Biology of the Cell. 17, 56–66 (2006).

18. C. E. Antal, J. D. Violin, M. T. Kunkel, S. Skovsø, A. C. Newton, Intramolecular conformational changes optimize protein kinase C signaling. Chemistry and Biology. 21, 459–469 (2014).

19. T. R. Baffi, A. A. N. Van, W. Zhao, G. B. Mills, A. C. Newton, Protein Kinase C Quality Control by Phosphatase PHLPP1 Unveils Loss-of-Function Mechanism in Cancer. Molecular Cell. 74, 378–392.e5 (2019).

20. I. Yabe, H. Sasaki, D. H. Chen, W. H. Raskind, T. D. Bird, I. Yamashita, S. Tsuji, S. Kikuchi, K. Tashiro, Spinocerebellar Ataxia Type 14 Caused by a Mutation in Protein Kinase C γ. Archives of Neurology. 60, 1749–1751 (2003).

21. I. Yamashita, H. Sasaki, I. Yabe, T. Fukazawa, S. Nogoshi, K. Komeichi, A. Takada, K. Shiraishi, Y. Takiyama, M. Nishizawa, J. Kaneko, H. Tanaka, S. Tsuji, K. Tashiro, A novel locus for dominant cerebellar ataxia (SCA14) maps to a 10.2-cM interval flanked by D19S206 and D19S605 on chromosome 19q13.4-qter. Annals of Neurology. 48, 156–163 (2000).

22. N. Adachi, T. Kobayashi, H. Takahashi, T. Kawasaki, Y. Shirai, T. Ueyama, T. Matsuda, T. Seki, N. Sakai, N. Saito, Enzymological analysis of mutant protein kinase Cγ causing spinocerebellar ataxia type 14 and dysfunction in Ca2+ homeostasis. Journal of Biological Chemistry. 283, 19854–19863 (2008).

23. M. M. K. Wong, S. D. Hoekstra, J. Vowles, L. M. Watson, G. Fuller, A. H. Németh, S. A. Cowley, O. Ansorge, K. Talbot, E. B. E. Becker, Neurodegeneration in SCA14 is associated with increased PKCγ kinase activity, mislocalization and aggregation. Acta neuropathologica communications. 6, 99 (2018).

24. T. Schmitz-Hübsch, S. Lux, P. Bauer, A. U. Brandt, E. Schlapakow, S. Greschus, M. Scheel, H. Gärtner, M. E. Kirlangic, V. Gras, D. Timmann, M. Synofzik, A. Giorgetti, P. Carloni, J. N. Shah, L. Schöls, U. Kopp, L. Bußenius, T. Oberwahrenbrock, H. Zimmermann, C. Pfueller, E. M. Kadas, M. Rönnefarth, A. S. Grosch, M. Endres, K. Amunts, F. Paul, S. Doss, M. Minnerop, Spinocerebellar ataxia type 14: refining clinicogenetic diagnosis in a rare adult-onset disorder. Annals of Clinical and Translational Neurology. 8, 774–789 (2021).

25. T. Shirafuji, H. Shimazaki, T. Miyagi, T. Ueyama, N. Adachi, S. Tanaka, I. Hide, N. Saito, N. Sakai, Spinocerebellar ataxia type 14 caused by a nonsense mutation in the PRKCG gene. Molecular and Cellular Neuroscience. 98, 46–53 (2019).

26. E. Shimobayashi, J. P. Kapfhammer, A New Mouse Model Related to SCA14 Carrying a Pseudosubstrate Domain Mutation in PKCγ Shows Perturbed Purkinje Cell Maturation and Ataxic Motor Behavior. Journal of Neuroscience. 41, 2053–2068 (2021).

27. J. Trzesniewski, S. Altmann, L. Jäger, J. P. Kapfhammer, Reduced Purkinje cell size is compatible with near normal morphology and function of the cerebellar cortex in a mouse model of spinocerebellar ataxia. Experimental Neurology. 311, 205–212 (2019).

28. K. Schrenk, J. P. Kapfhammer, F. Metzger, Altered dendritic development of cerebellar Purkinje cells in slice cultures from protein kinase Cγ-deficient mice. Neuroscience. 110, 675–689 (2002).

29. A. M. Ghoumari, R. Wehrlé, C. I. De Zeeuw, C. Sotelo, I. Dusart, Inhibition of Protein Kinase C Prevents Purkinje Cell Death but Does Not Affect Axonal Regeneration. Journal of Neuroscience. 22, 3531–3542 (2002).

30. D. S. Verbeek, J. Goedhart, L. Bruinsma, R. J. Sinke, E. A. Reits, PKCγ mutations in spinocerebellar ataxia type 14 affect C1 domain accessibility and kinase activity leading to aberrant MAPK signaling. Journal of Cell Science. 121, 2339–2349 (2008).

31. D. S. Verbeek, M. A. Knight, G. G. Harmison, K. H. Fischbeck, B. W. Howell, Protein kinase C gamma mutations in spinocerebellar ataxia 14 increase kinase activity and alter membrane targeting. Brain. 128, 436–442 (2005).

32. H. Takahashi, N. Adachi, T. Shirafuji, S. Danno, T. Ueyama, M. Vendruscolo, A. N. Shuvaev, T. Sugimoto, T. Seki, D. Hamada, K. Irie, H. Hirai, N. Sakai, N. Saito, Identification and characterization of PKCγ, a kinase associated with SCA14, as an amyloidogenic protein. Human Molecular Genetics. 24, 525–539 (2015).

33. A. Nakazono, N. Adachi, H. Takahashi, T. Seki, D. Hamada, T. Ueyama, N. Sakai, N. Saito, Pharmacological induction of heat shock proteins ameliorates toxicity of mutant PKCɣ in spinocerebellar ataxia type 14. Journal of Biological Chemistry. 293, 14758– 14774 (2018).

34. T. Seki, T. Shimahara, K. Yamamoto, N. Abe, T. Amano, N. Adachi, H. Takahashi, K. Kashiwagi, N. Saito, N. Sakai, Mutant γPKC found in spinocerebellar ataxia type 14 induces aggregate-independent maldevelopment of dendrites in primary cultured Purkinje cells. Neurobiology of Disease. 33, 260–273 (2009).

35. M. G. Kazanietz, N. E. Lewin, J. D. Bruns, P. M. Blumberg, Characterization of the cysteine-rich region of the Caenorhabditis elegans protein Unc-13 as a high affinity phorbol ester receptor. Analysis of ligand- binding interactions, lipid cofactor requirements, and inhibitor sensitivity. Journal of Biological Chemistry. 270, 10777– 10783 (1995).

36. B. L. Ross, B. Tenner, M. L. Markwardt, A. Zviman, G. Shi, J. P. Kerr, N. E. Snell, J. J. McFarland, J. R. Mauban, C. W. Ward, M. A. Rizzo, J. Zhang, Single-color, ratiometric biosensors for detecting signaling activities in live cells. eLife. 7 (2018), doi:10.7554/elife.35458.

37. L. L. Gallegos, M. T. Kunkel, A. C. Newton, Targeting protein kinase C activity reporter to discrete intracellular regions reveals spatiotemporal differences in agonist-dependent signaling. Journal of Biological Chemistry. 281, 30947–30956 (2006).

38. J. D. Violin, J. Zhang, R. Y. Tsien, A. C. Newton, A genetically encoded fluorescent reporter reveals oscillatory phosphorylation by protein kinase C. Journal of Cell Biology. 161, 899–909 (2003).

39. L. M. Keranen, E. M. Dutil, A. C. Newton, Protein kinase C is regulated in vivo by three functionally distinct phosphorylations. Current Biology. 5, 1394–1403 (1995).

40. J. Jezierska, J. Goedhart, H. H. Kampinga, E. A. Reits, D. S. Verbeek, SCA14 mutation V138E leads to partly unfolded PKCγ associated with an exposed C-terminus, altered kinetics, phosphorylation and enhanced insolubilization. Journal of Neurochemistry. 128, 741–751 (2014).

41. D. J. Burns, R. M. Bell, Protein kinase C contains two phorbol ester binding domains. Journal of Biological Chemistry. 266, 18330–18338 (1991).

42. Y. Xiao, T. H. Hsiao, U. Suresh, H. I. H. Chen, X. Wu, S. E. Wolf, Y. Chen, A novel significance score for gene selection and ranking. Bioinformatics. 30, 801–807 (2014).

43. S. Guidato, L. H. Tsai, J. Woodgett, C. C. J. Miller, Differential cellular phosphorylation of neurofilament heavy side-arms by glycogen synthase kinase-3 and cydin-dependent kinase-5. Journal of Neurochemistry. 66, 1698–1706 (1996).

44. S. Lee, H. C. Pant, T. B. Shea, Divergent and convergent roles for kinases and phosphatases in neurofilament dynamics. Journal of Cell Science. 127, 4064–4077 (2014).

45. T. M. Thornton, G. Pedraza-Alva, B. Deng, C. D. Wood, A. Aronshtam, J. L. Clements, G. Sabio, R. J. Davis, D. E. Matthews, B. Doble, M. Rincon, Phosphorylation by p38 MAPK as an alternative pathway for GSK3β inactivation. Science. 320, 667–670 (2008).

46. D. W. Huang, B. T. Sherman, R. A. Lempicki, Systematic and integrative analysis of large gene lists using DAVID bioinformatics resources. Nature Protocols. 4, 44–57 (2009).

47. D. W. Huang, B. T. Sherman, R. A. Lempicki, Bioinformatics enrichment tools: Paths toward the comprehensive functional analysis of large gene lists. Nucleic Acids Research. 37, 1–13 (2009).

48. C. E. Antal, A. M. Hudson, E. Kang, C. Zanca, C. Wirth, N. L. Stephenson, E. W. Trotter, L. L. Gallegos, C. J. Miller, F. B. Furnari, T. Hunter, J. Brognard, A. C. Newton, Cancer-associated protein kinase C mutations reveal kinase’s role as tumor suppressor. Cell. 160, 489–502 (2015).

49. E. Cerami, J. Gao, U. Dogrusoz, B. E. Gross, S. O. Sumer, B. A. Aksoy, A. Jacobsen, C. J. Byrne, M. L. Heuer, E. Larsson, Y. Antipin, B. Reva, A. P. Goldberg, C. Sander, N. Schultz, The cBio Cancer Genomics Portal: An open platform for exploring multidimensional cancer genomics data. Cancer Discovery. 2, 401–404 (2012).

50. J. Gao, B. A. Aksoy, U. Dogrusoz, G. Dresdner, B. Gross, S. O. Sumer, Y. Sun, A. Jacobsen, R. Sinha, E. Larsson, E. Cerami, C. Sander, N. Schultz, Integrative analysis of complex cancer genomics and clinical profiles using the cBioPortal. Science Signaling. 6, 1–1 (2013).

51. R. L. Grossman, A. P. Heath, V. Ferretti, H. E. Varmus, D. R. Lowy, W. A. Kibbe, L. M. Staudt, Toward a Shared Vision for Cancer Genomic Data. New England Journal of Medicine. 375, 1109–1112 (2016).

52. M. H. M. Vlak, R. J. Sinke, G. M. Rabelink, B. P. H. Kremer, B. P. C. van de Warrenburg, Novel PRKCG/SCA14 mutation in a Dutch spinocerebellar ataxia family: Expanding the phenotype. Movement Disorders. 21, 1025–1028 (2006).

53. C. Ganos, S. Zittel, M. Minnerop, O. Schunke, C. Heinbokel, C. Gerloff, C. Zühlke, P. Bauer, T. Klockgether, A. Münchau, T. Bäumer, Clinical and neurophysiological profile of four German families with spinocerebellar ataxia type 14. Cerebellum. 13, 89–96 (2014).

54. V. Chelban, S. Wiethoff, B. K. Fabian-Jessing, N. A. Haridy, A. Khan, S. Efthymiou, E. B. E. Becker, E. O’Connor, J. Hersheson, K. Newland, A. T. Hojland, P. A. Gregersen, S. G. Lindquist, M. B. Petersen, J. E. Nielsen, M. Nielsen, N. W. Wood, P. Giunti, H. Houlden, Genotype-phenotype correlations, dystonia and disease progression in spinocerebellar ataxia type 14. Movement Disorders. 33, 1119–1129 (2018).

55. G. Stevanin, V. Hahn, E. Lohmann, N. Bouslam, M. Gouttard, C. Soumphonphakdy, M. L. Welter, E. Ollagnon-Roman, A. Lemainque, M. Ruberg, A. Brice, A. Durr, Mutation in the catalytic domain of protein kinase C γ and extension of the phenotype associated with spinocerebellar ataxia type 14. Archives of Neurology. 61, 1242–1248 (2004).

56. S. Klebe, A. Durr, A. Rentschler, V. Hahn-Barma, M. Abele, N. Bouslam, L. Schöls, P. Jedynak, S. Forlani, E. Denis, C. Dussert, Y. Agid, P. Bauer, C. Globas, U. Wüllner, A. Brice, O. Riess, G. Stevanin, New mutations in protein kinase Cγ associated with spinocerebellar ataxia type 14. Annals of Neurology. 58, 720–729 (2005).

57. D. Chen, P. Cimino, L. Ranum, H. Zoghbi, I. Yabe, L. Schut, R. Margolis, H. Lipe, A. Feleke, M. Matsushita, J. Wolff, C. Morgan, D. Lau, M. Fernandez, H. Sasaki, W. Raskind, T. Bird, The clinical and genetic spectrum of spinocerebellar ataxia 14. Neurology. 64, 1258–60 (2005).

58. N. Kannan, N. Haste, S. S. Taylor, A. F. Neuwald, The hallmark of AGC kinase functional divergence is its C-terminal tail, a cis-acting regulatory module. Proceedings of the National Academy of Sciences of the United States of America. 105 (2008), p. 9130.

59. T. A. Leonard, B. Róycki, L. F. Saidi, G. Hummer, J. H. Hurley, Crystal structure and allosteric activation of protein kinase C βiI. Cell. 144, 55–66 (2011).

60. G. Hansra, P. Garcia-Paramio, C. Prevostel, R. D. H. Whelan, F. Bornancin, P. J. Parker, Multisite dephosphorylation and desensitization of conventional protein kinase C isotypes. Biochemical Journal. 342, 337–344 (1999).

61. D. Chen, C. Gould, R. Garza, T. Gao, R. Y. Hampton, A. C. Newton, Amplitude control of protein kinase C by RINCK, a novel E3 ubiquitin ligase. Journal of Biological Chemistry. 282, 33776–33787 (2007).

62. O. V. Leontieva, J. D. Black, Identification of Two Distinct Pathways of Protein Kinase Cα Down-regulation in Intestinal Epithelial Cells. Journal of Biological Chemistry. 279, 5788–5801 (2004).

63. U. Kirchhefer, A. Heinick, S. König, T. Kristensen, F. U. Müller, M. D. Seidl, P. Boknik, Protein phosphatase 2A is regulated by protein kinase Cα (PKCα)-dependent phosphorylation of its targeting subunit B56α at Ser41. Journal of Biological Chemistry. 289, 163–176 (2014).

64. J. H. Ahn, T. McAvoy, S. V Rakhilin, A. Nishi, P. Greengard, A. C. Nairn, Protein kinase A activates protein phosphatase 2A by phosphorylation of the B56δ subunit. Proceedings of the National Academy of Sciences of the United States of America. 104, 2979–2984 (2007).

65. J. H. Ahn, Y. Kim, H. S. Kim, P. Greengard, A. C. Nairn, Protein kinase C-dependent dephosphorylation of tyrosine hydroxylase requires the B56δ heterotrimeric form of protein phosphatase 2A. PLoS ONE. 6 (2011), doi:10.1371/journal.pone.0026292.

66. A. Yuan, M. V. Rao, Veeranna, R. A. Nixon, Neurofilaments at a glance. Journal of Cell Science. 125, 3257–3263 (2012).

67. N. Goode, K. Hughes, J. R. Woodgett, P. J. Parker, Differential regulation of glycogen synthase kinase-3β by protein kinase C isotypes. Journal of Biological Chemistry. 267, 16878–16882 (1992).

68. E. A. R. Nibbeling, A. Duarri, C. C. Verschuuren-Bemelmans, M. R. Fokkens, J. M. Karjalainen, C. J. L. M. Smeets, J. J. De Boer-Bergsma, G. Van Der Vries, D. Dooijes, G. B. Bampi, C. Van Diemen, E. Brunt, E. Ippel, B. Kremer, M. Vlak, N. Adir, C. Wijmenga, B. P. C. Van De Warrenburg, L. Franke, R. J. Sinke, D. S. Verbeek, Exome sequencing and network analysis identifies shared mechanisms underlying spinocerebellar ataxia. Brain. 140, 2860–2878 (2017).

69. R. Tsumagari, S. Kakizawa, S. Kikunaga, Y. Fujihara, S. Ueda, M. Yamanoue, N. Saito, M. Ikawa, Y. Shirai, DGKγ Knock-Out Mice Show Impairments in Cerebellar Motor Coordination, LTD, and the Dendritic Development of Purkinje Cells through the Activation of PKCγ. eNeuro. 7 (2020), doi:10.1523/ENEURO.0319-19.2020.

70. M. Leitges, J. Kovac, M. Plomann, D. J. Linden, A unique PDZ ligand in PKCα confers induction of cerebellar long-term synaptic depression. Neuron. 44, 585–594 (2004).

71. C. Chen, M. Kano, A. Abeliovich, L. Chen, S. Bao, J. J. Kim, K. Hashimoto, R. F. Thompson, S. Tonegawa, Impaired motor coordination correlates with persistent multiple climbing fiber innervation in PKCγ mutant mice. Cell. 83, 1233–1242 (1995).

72. A. N. Shuvaev, H. Horiuchi, T. Seki, H. Goenawan, T. Irie, A. Iizuka, N. Sakai, H. Hirai, Mutant PKCγ in spinocerebellar ataxia type 14 disrupts synapse elimination and long-term depression in purkinje cells in vivo. Journal of Neuroscience. 31, 14324–14334 (2011).

73. J. Van Baal, J. De Widt, N. Divecha, W. J. Van Blitterswijk, Translocation of diacylglycerol kinase θ from cytosol to plasma membrane in response to activation of G protein-coupled receptors and protein kinase C. Journal of Biological Chemistry. 280, 9870–9878 (2005).

74. L. L. Gallegos, A. C. Newton, Genetically encoded fluorescent reporters to visualize protein kinase C activation in live cells. Methods in Molecular Biology. 756, 295–310 (2011).

75. L. M. Keranen, A. C. Newton, Ca2+ differentially regulates conventional protein kinase Cs’ membrane interaction and activation. Journal of Biological Chemistry. 272, 25959– 25967 (1997).

76. J. D. Lapek, M. K. Lewinski, J. M. Wozniak, J. Guatelli, D. J. Gonzalez, Quantitative temporal viromics of an inducible HIV-1 model yields insight to global host targets and phospho-dynamics associated with protein Vpr. Molecular and Cellular Proteomics. 16, 1447–1461 (2017).

77. E. F. Pettersen, T. D. Goddard, C. C. Huang, G. S. Couch, D. M. Greenblatt, E. C. Meng, T. E. Ferrin, UCSF Chimera - A visualization system for exploratory research and analysis. Journal of Computational Chemistry. 25, 1605–1612 (2004).

78. A. Šali, T. L. Blundell, Comparative Protein Modelling by Satisfaction of Spatial Restraints. Journal of Molecular Biology. 234 (1993), pp. 779–815.

79. D. Kozakov, D. R. Hall, B. Xia, K. A. Porter, D. Padhorny, C. Yueh, D. Beglov, S. Vajda, The ClusPro web server for protein-protein docking. Nature Protocols. 12, 255–278 (2017).

80. S. El-Gebali, J. Mistry, A. Bateman, S. R. Eddy, A. Luciani, S. C. Potter, M. Qureshi, L. J. Richardson, G. A. Salazar, A. Smart, E. L. L. Sonnhammer, L. Hirsh, L. Paladin, D. Piovesan, S. C. E. Tosatto, R. D. Finn, The Pfam protein families database in 2019. Nucleic Acids Research. 47, D427–D432 (2019).

